# Circular RNA signatures of human healing and non-healing wounds

**DOI:** 10.1101/2021.11.23.469681

**Authors:** Maria A. Toma, Zhuang Liu, Qizhang Wang, Letian Zhang, Dongqing Li, Pehr Sommar, Ning Xu Landén

## Abstract

**Background:** Although the widespread expression of circular RNAs (circRNAs) has only been recognized recently, increasing evidence has suggested their important roles in health and disease. To identify clinically relevant circRNAs with potential for wound diagnosis and therapy, an in-depth characterization of circRNA expression in human healing and non-healing wounds is a prerequisite that has not been attained yet.

**Methods:** We collected wound-edge biopsies through the healing process of healthy donors and in chronic non-healing venous ulcers (VU). Paired total RNA- and small RNA-sequencing were performed to profile circRNAs, protein-coding mRNAs, and microRNA expression. We analyzed the co-expression relationship between circRNAs and mRNAs with weighted correlation network analysis (WGCNA) and constructed circRNA-microRNA-mRNA networks. For the circRNAs surfaced in the in-silico analysis, after validating their expression with RT-PCR and sequencing, we silenced hsa-CHST15_0003 and hsa-TNFRSF21_0001 expression in keratinocytes with siRNAs and studied their function with transcriptomic profiling and live-cell monitoring.

**Results:** Our study unravels the dynamically changed expression patterns of circRNAs during human skin wound healing and their abnormal expression signature in VU, which are presented as a searchable web resource (http://130.229.28.87/shiny/circRNA_wholebiopsy-shinyApp/). *In silico* analysis deciphers the circRNA-miRNAs-mRNA networks specific to the inflammatory and proliferative phases of wound repair and VU, the biological processes that circRNAs are involved, and the circRNAs that could act as miRNAs sponge in human wounds. Importantly, we found that hsa-CHST15_0003 and hsa-TNFRSF21_0001, two circRNAs upregulated in VU, hampered keratinocyte migration while promoting proliferation through modulating gene networks underpinning these cellular processes.

**Conclusion:** By integrating circRNA, mRNA, and miRNA expression profiles in a unique collection of clinical samples, we identify the circRNAs that are relevant to human wound healing physiology and pathology. This study paves the way to decipher the functional significance of circRNAs in tissue repair.

## Background

Skin wound healing is an intricate process that requires the timely cooperation of multiple cell types and occurs in three sequential but overlapping phases, i.e., inflammation, growth, and remodeling (1). Venous ulcers (VU) are a type of chronic non-healing wounds and represent the most common form of leg ulcers (2). VUs occur mainly in patients with chronic venous disease and venous hypertension (3) and represent an important medical problem and an increasing burden for healthcare and society. It is estimated that 5-8% of the world population suffer from venous disease and 20% of these patients develop VUs (4). Due to the complex wound environment, many patients fail to respond to the standard VU therapy. There is thus a need for a better understanding of the molecular events that characterize VU pathology.

Circular RNA (circRNA) is a novel class of RNA, which widespread expression in eukaryotes has only been recognized lately (5). They have a covalent contiguous loop structure without 5’ and 3’ ends and are mainly formed by ‘back-splicing’ of pre-mRNA, in which a downstream 5’splice site is joined to an upstream 3’splice site in a reverse order (5). To date, most studies about circRNAs provide evidence for their role as competing endogenous RNAs that sponge microRNAs (miRNAs) to rescue the expression of the miRNA target genes (6). Also, some circRNAs have been shown to interact with proteins and act as decoys, scaffolds, or recruiting partners (5). A growing number of circRNAs has been implicated in the regulation of skin biology, including epidermal differentiation (7), melanogenesis (8), and aging (9), as well as skin diseases, such as psoriasis, atopic dermatitis, and skin cancers [reviewed in (10)]. However, their functional characterization in skin wound healing is in its infancy, and to our knowledge, studies addressing the roles of circRNAs in VUs are still lacking to date.

To fill up these knowledge gaps, we performed paired total RNA-sequencing (seq) and small RNA-seq in the tissue samples of human skin, acute wounds at inflammatory and proliferative phases, and VUs. This enables us to capture the genome-wide transcriptomic changes, including circRNAs, mRNAs, and miRNAs, during skin wound healing and the pathological gene signature of the non-healing VUs. Here, we focused on the circRNAs to characterize their global expression pattern and identify the differentially expressed ones in human wounds. Furthermore, we inferred the functions of circRNAs by analyzing the *in vivo* co-expression networks with circRNAs and mRNAs and highlighted the circRNAs that may act via sponging miRNAs in wound repair. The wound-associated circRNAs were experimentally validated for their expression and function. In particularly, hsa-CHST15_0003 and hsa-TNFRSF21_0001 were found to regulate epidermal keratinocyte migration and growth. We presented this human wound circRNA expression landscape at a browsable web portal (http://130.229.28.87/shiny/circRNA_wholebiopsy-shinyApp/), to prime further search of clinical-relevant circRNAs promoting tissue repair and regeneration.

## Materials and methods

### Human skin wound specimens

To investigate circRNA expression profiles *in vivo,* human skin and wound biopsies were obtained from 10 healthy volunteers and 12 chronic VU patients. Detailed donor and patient information are presented in **Table S1-3.** The exclusion criteria for healthy donors were diabetes, skin disease, unstable heart disease, infections, bleeding disorder, immune suppression, and any ongoing medical treatments. On the skin of each healthy donor, two excisional wounds were created using a 4□mm punch and the excised skin from these surgical wounds was saved as intact skin control. The wound edges were collected using a 6 mm punch one day (Wound1) and seven days later (Wound7) (**Figure 1a**). Wound-edge samples were collected from patients with non-healing VU that, despite conventional therapy, persisted for more than four months (**Table S3**). Biopsies were taken from the non-healing edge of VUs using a 4□mm punch (**Figure 1a**). Local lidocaine injection was used for anesthesia while sampling. Written informed consent was obtained from all donors for the collection and use of clinical samples. The study was approved by the Stockholm Regional Ethics Committee and conducted according to the Declaration of Helsinki’s principles.

**Figure 1.**
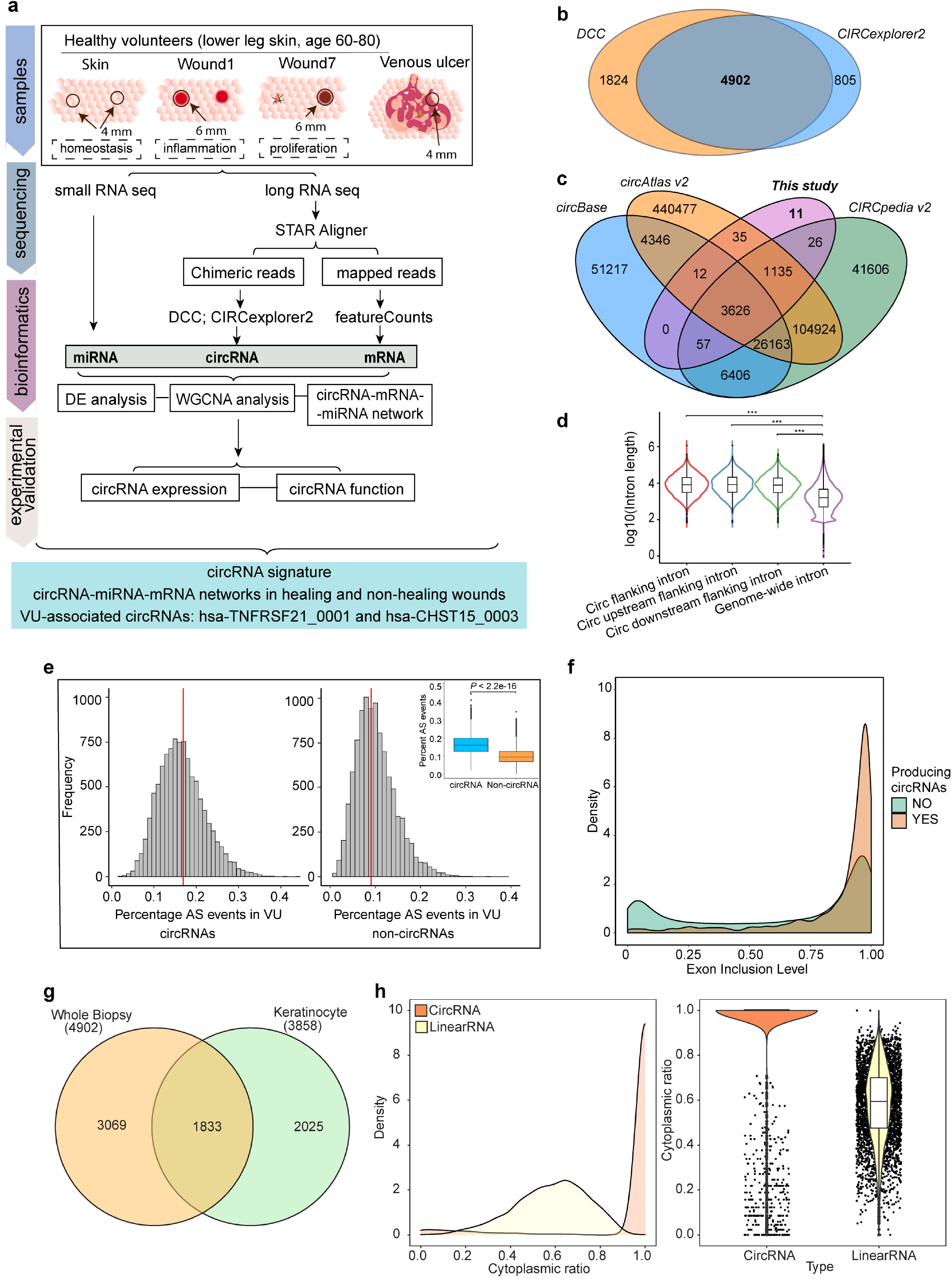
Profiling of circular RNA expression in human skin wounds. (**a**) Flowchart of the design and outcome of this study. (**b**) Venn diagram shows the number of circRNAs detected by DCC and CIRCexplorer2 (BSJ reads ≥ 2 in at least 2 samples). (**c**) Venn diagram depicts the number of annotated circRNAs by using databases circBase, circAtlas version 2, and CIRCpedia version 2. (**d**) A violin plot shows the length of flanking introns of circRNA-generating exons vs. genome-wide exons. (**e**) Histograms show the frequency of alternative splicing (AS) event percentage in circRNA-forming exons (left) and non-circRNA-forming exons (right) after 10000 random samplings corrected for intron lengths and gene expression levels in VU. (**f**) Density plot of exon inclusion level for circRNA-forming exons and non-circRNA-forming exons. (**g**) Venn diagram depicts the numbers of circRNAs detected in full-depth wound biopsies and human primary keratinocytes by RNA-seq. (**h**) Density (left) and violin plots (right) of cytoplasmic ratios of circRNAs and linearRNAs in keratinocytes. (**d, e**) Mann-Whitney-Wilcoxon Test, *\*\*\*P* < 0.001.

### Cell culture

Primary human epidermal keratinocytes (Cascade Biologics, Portland, OR) were cultured in EpiLife medium supplemented with 10% Human Keratinocyte Growth Supplement (HKGS) and 1% penicillin/streptomycin. Primary human dermal fibroblasts, adult (Cascade Biologics) were cultured in Medium 106 supplemented with 10% Low Serum Growth Supplement (LSGS) and 1% penicillin/streptomycin. Cells were grown at 37°C in 5% CO_2_ (ThermoFisher Scientific, Carlsbad, CA).

### Cell fractionation

Cytoplasm and nucleus of keratinocytes were separated by using Nuclear Extract Kit (Active Motif, Inc., Carlsbad, CA, USA) following the manufacturer’s instructions. Total RNA was extracted from these fractions using miRNeasy Kit (Qiagen).

### RNA-seq library preparation and sequencing

Total RNAs were isolated from whole biopsies and isolated cytoplasm and nucleus of keratinocytes, i.e., the skin, Wound1, Wound7, and VU (n=5/ each group), cytoplasm and nucleus (n=1/each group), by using the miRNeasy Mini kit (Qiagen, Hilden, Germany) and prepared for library construction. First, the ribosomal RNA (rRNA) was removed using the Epicentre Ribo-zero® rRNA Removal Kit (Epicentre) with a total amount of 2 ug RNA as an input for each library. Second, strand-specific RNA-seq libraries were constructed by using the NEB Next® UltraTM Directional RNA Library Prep Kit for Illumina® (NEB) according to the manufacturer’s instructions. Finally, the libraries were sequenced on the Illumina Hiseq 4000 platform (Illumina, Inc.) by using 150 bp paired-end reads.

### CircRNA identification and analysis

#### CircRNA identification

Raw reads were first filtered by using the Trimmomatic v0.36 package (11) to remove adaptor sequences and low-quality bases. Clean reads were then mapped to human reference genome (GRCh38.p12) with the GENCODE genes annotation (version 31) using STAR v2.7.1 (12). Gene expression of mRNAs was determined by counting uniquely mapped fragments to exons using featureCount package (13). Raw counts for each mRNA were normalized to FPKM (fragments per kilobase of transcript, per million mapped reads). The unmapped chimeric reads were used for circRNA identification using CIRCexplorer2 (14) and DCC (15). In addition, the cognate linear RNA expression was determined from the average read depth of linear-spliced reads close to the back-splice junction sites at both flanking exons by using the DCC package (15). CircRNAs were filtered by at least two back-spliced junction reads in a minimum of two samples. And only circRNAs overlapped by two tools were kept for the following analysis. Expression of circRNAs was normalized to FPM (fragments mapped to back-splicing junctions per million mapped fragments) as previously described (14). CircRNAs were annotated based on chromosomal location and the overlap with three circRNA databases: circAtlas v2 (16), circBase (17), and CIRCpedia v2 (18). The circRNA nomenclature used in this paper is based on circAtlas v2. In addition, the cytoplasmic ratios of circRNA or linearRNA of keratinocytes were calculated as previously described (19): Cytoplasmic ratio = Cytoplasmic normalized expression / (Cytoplasmic normalized expression + Nuclear normalized expression).

#### Alternative splicing (AS) analysis

Pairwise differential AS events within four groups were identified using rMATS.4.0.2 with following option ‘-readLength 150 -cstat 0.0001 -t paired -libType fr-firststrand’ (20). AS events in each group were then filtered by at least two inclusion junction reads and two skipping junction reads in two independent samples. To determine whether the number of AS events located in exons generating circRNAs was significantly higher than that of exons without circRNAs, we randomly selected an equal number of exons not generating circRNAs but with the same range of mRNA expression levels and intron lengths. The simulation analysis was repeated 1000 times using the custom script.

#### Differential expression (DE) analysis

Prior to DE analysis, principle component analysis (PCA) was carried out on the transformed circRNA expression data using the plotPCA function. Differential circRNA and cognate linear RNA analysis were then performed by using DESeq2 package (21). P-values calculated from the Wald test were adjusted by performing Benjamini-Hochberg (BH) multiple testing to estimate false discovery rate (FDR). The differentially expressed circular RNAs or cognate linear RNAs were defined as fold change greater than 2 and FDR < 0.05. Plots were drawn by using ggplot2 (22) and ComplexHeatmap packages (23).

### Co-expression network analysis of circRNAs-mRNAs

Weighted gene co-expression network analysis (WGCNA) was performed on the combined circRNA and mRNA expression data. CircRNAs expressed in at least half of the twenty samples and mRNAs differentially expressed (FDR < 0.05 and |log2(fold change)| ≥ 1) in six pairwise comparisons were included. The co-expression network was constructed with normalized expression of 2187 circRNAs and 8347 DE mRNAs using the one-step blockwiseModules algorithm in the WGCNA R package (24). Eighteen modules were identified with the following parameters: power = 16, networkType = “Signed”, corFnc = “bicor”, minModulesIZE = 30, mergeCutHeight = 0.25. The optimal threshold power of 16 was determined according to the fit of scale-free topological network (**Figure S4a**). CircRNAs-mRNAs were assigned to a module based on their correlations to the module eigengenes (ME), which represents the first principal component of all circRNAs and mRNAs in a module, and the significant correlations were filtered by P adjusted values using the BH method. We performed gene ontology (GO) enrichment analysis for each condition with the clusterProfiler package (25) by using the mRNAs in significant modules. GO terms of biological process (BP) were ranked based on gene numbers in each GO term in ascending order and filtered by FDR < 0.01. Furthermore, module preservation analysis were carried out based on the same twenty samples as reference and test datasets by permutating 200 times using the modulePreservation function from the WGCNA R package (24).

### Identification of circRNA-miRNA-mRNA interactions

For the circRNAs in the WGCNA modules significantly associated with each condition, we constructed circRNA-miRNA-mRNA networks (**Figure 4a**). First, miRNA normalized expression data were obtained from small RNA-sequencing of the same sample cohort (unpublished) and 562 miRNAs that were expressed in at least 10 of the 20 samples were included in this analysis. Second, to determine the circRNA-miRNA interactions, we searched for miRNA target sites within circRNA sequences by using the RNA22 v2 (26) and TargetScan v7.2 (27) packages, and only kept the circRNA-miRNA interactions commonly predicted by both tools. Third, we inferred the miRNA-mRNA interactions with both TargetScan (27) and miRDB (28) databases by using the miRWalk tool (29), and only the mRNAs detected in more than half of our samples were included in this analysis. Fourth, we connected circRNAs and mRNAs through the common miRNAs they could bind to. Finally, we performed the Spearman rank correlation analysis in pairwise connections among circRNAs, miRNAs, and mRNAs using their normalized expression profiles. Only circRNA-miRNA and miRNA-mRNA interactions with significantly negative correlations, and circRNA-mRNA interactions with positive correlations (Spearman’s P value < 0.05) were reserved in the final networks.

### Experimental validation of candidate circRNAs

#### Cell transfection

Third passage keratinocytes were seeded in 24-well plates. At 60-70% confluence, cells were transfected with 20nM siRNA against hsa-CHST15_0003 or hsa-TNFRSF21_0001 or a non-targeting siRNA control (GE Healthcare Dharmacon, Inc., Lafayette, CO, USA) using Lipofectamine 3000 (ThermoFisher Scientific). After 24 hours, cells were harvested for RNA extraction.

#### RNA extraction

Tissue biopsies were homogenized using TissueLyser LT (Qiagen) before RNA extraction. Total RNA was extracted from human tissue and cells using the miRNeasy Mini kit (Qiagen) or TRIzol reagent (ThermoFisher Scientific).

#### RNase R digestion

To assess whether candidate circRNAs are resistant to degradation by RNase R, 3 ug of total RNA from cultured keratinocytes or fibroblasts was treated without (control) or with 10 units of RNase R in 10X RNase R reaction buffer (LGC Biosearch Technologies, Hoddesdon, UK). Treatment was conducted at 37°C for 12 minutes, followed by RNase R inactivation at 70°C for 10 min. The digested RNA and the control RNA were then converted to cDNA by using RevertAid First Strand cDNA Synthesis Kit (ThermoFisher Scientific) with random primers.

#### RT-PCR, DNA electrophoresis, and Sanger sequencing

To verify the authenticity of the circRNAs identified by *in silico* analysis of the RNA-seq data, we used PCR with outward-facing primers to detect head-to-tail splice sites of 15 circRNA candidates. Fifty ng cDNA obtained from keratinocytes or fibroblasts was amplified with divergent primers, which span over circRNA-forming back-splice junction (**Table S6**), by using PCR Master Mix (2X) (ThermoFisher Scientific) and following the recommended thermal conditions for 33 cycles. The PCR products were then analyzed by DNA electrophoresis using E-Gel™ Agarose Gels with SYBR™ Safe DNA Gel Stain, 2% (Invitrogen, Waltham, MA, USA). The bands found at expected sizes were cut out and the DNA was purified by using QIAquick Gel Extraction Kit (Qiagen). Purified DNA was then analyzed by Sanger sequencing using ABI 3730 PRISM® DNA Analyzer at the KIgene core facility at Karolinska Institutet, Stockholm, Sweden.

#### qRT-PCR

To quantify circRNA expression, we performed quantitative Real-time PCR (qRT-PCR) assays by using SYBR™ Green PCR Master Mix (Applied Biosystems, Waltham, MA). These experiments were carried out with the QuantStudio 6 and 7 Flex Real-Time PCR Systems (Applied Biosystems). Expression of circRNAs was calculated relative to the housekeeping genes *18S* or *ACTB*. Sequences of the primers were depicted in **Table S6**.

### Functional analysis of circRNAs

#### Gene expression microarray

Gene expression profiling of keratinocytes transfected with either siRNA control (*n* = 3) or si-hsa-CHST15_0003 (*n*□=□3) or si-hsa-TNFRSF21_0001 (*n*□=□3) for 24□hours was carried out by using human Clariom™ Sassay (ThermoFisher Scientific) at the Bioinformatics and Expression Analysis (BEA) core facility at Karolinska Institutet. In brief, total RNA was extracted using TRIzol, and RNA quality and quantity were determined using Agilent 2200 Tapestation with RNA ScreenTape and Nanodrop 1000. One hundred fifty nanograms of total RNA was used to prepare cDNA following the GeneChip WT PLUS Reagent Kit labeling protocol. Standardized array processing procedures recommended by Affymetrix, including hybridization, fluidics processing, and scanning, were used. Expression data were analyzed by using Transcriptome Analysis Console 4.0. The data discussed herein have been deposited in the NCBI’s Gene Expression Omnibus database under accession number GSE188939.

#### Gene set enrichment analysis (GSEA) analysis

Using the fgsea R package, we performed GSEA for biological process (BP), Kyoto Encyclopedia of Genes and Genomes (KEGG) pathway, and hallmark from Molecular Signatures Database (MSigDB) (http://www.gsea-msigdb.org/) with a ranked fold change list of all the genes detected in the microarray (30) (31). Using the STRING App and Cytoscape (version 3.7.2), we generated functional protein association networks for the genes regulated by hsa-CHST15_0003 and hsa-TNFRSF21_0001. CentiScape 2.2 App was used to characterize topological features of each node. Central node genes with high *Degree* values, which depict the number of links of each node, were chosen for validation by qRT-PCR.

#### Analysis of cell motility and cell growth with IncuCyte Live-cell imaging

Keratinocytes with hsa-CHST15_0003 or hsa-TNFRSF21_0001 knockdown were analyzed for cell migration or growth with IncuCyte™ live-cell imaging (Essen BioScience, Ann Arbor, MI).

In migration assays, cells were seeded onto a 96-well microtiter IncuCyte® ImageLock Plate (Essen BioScience) pre-coated with rat tail Collagen type I (Gibco™). When cells were at full confluence, standardized wounds were created by using Essen® 96-pin WoundMaker™ (Essen BioScience). The complete medium was changed for EpiLife medium without any supplements to avoid confounding effects caused by cell proliferation. The assay plates were incubated in the IncuCyte and scanned every 2□hours for 24□hours. The data were analyzed with the integrated metrics in the IncuCyte ZOOM software package and presented as the migration rate.

For cell growth monitoring, transfected cells were seeded at low density 1000 cells/well in 96-well microtiter IncuCyte® ImageLock Plate (Essen BioScience) pre-coated with rat tail Collagen type I (Gibco™). After 24Lhours, the assay plates were incubated in the IncuCyte and scanned every 2□hours for 60□hours. The cell confluence was analyzed with the IncuCyte ZOOM software package and expressed as Phase Count Object (PCO). The results were presented as the proliferation rate = (PCO_timepoint x_−PCO_timepoint 0_)x100/PCO_timepoint 0_.

### Statistical analysis

Data analysis was performed by using R and GraphPad 8.4.0 (GraphPad Software). Normality and distribution of data was checked with Shapiro-Wilk test (p < 0.05 indicated data that did not pass the normality test). Comparison between two groups was performed with unpaired two-tailed Student’s t-test (parametric) or Mann-Whitney U test (non-parametric, unpaired). Comparison between more than two groups that contained paired data (matched samples or repeated measures) was done with RM one-way ANOVA and Tukey’s multiple comparisons test (parametric data) or Friedman test and Dunn’s multiple comparisons test (non-parametric data). Comparison between more than two groups with unpaired data was performed with Ordinary one-way ANOVA and Dunnett’s multiple comparisons test (parametric data) or Kruskal-Wallis and Dunn’s multiple comparisons test (non-parametric data). p value < 0.05 was considered statistically significant.

## Results

### Circular RNA expression landscape in human skin wounds

To prime the complex gene expression regulation network underlying the wound healing process of human skin, we profiled the short and long RNA expression in the skin and acute wounds of five healthy donors with standardized excisional wounds and in the VUs from five patients (**Figure 1a**). The acute wound tissues were collected one and seven days after wounding, which allowed us to map transcriptomic changes during human skin wound repair at the inflammatory phase (Wound1) and the proliferative phase (Wound7). Moreover, the healthy donors (n=10) and VU patients (n=12) enrolled in this study were well matched in age group, sex, ethnicity, and anatomical location of the wounds, which set a good basis to compare the healing and chronic non-healing human wounds (**Table S1-S3**). Considering most knowledge in wound biology has been gained from non-human models, here we have a unique opportunity to probe the healing mechanisms of human skin, and to gain a deeper understanding about VU pathology, a complex disease that cannot be fully recapitulated in animal models (32).

The current study focused on the identification of circRNAs that may play a physiological or pathological role in wound healing. For this, we sequenced the ribosomal (r)RNA-depleted total RNA from the above clinical samples. After filtering the raw data with stringent criteria (**Table S4**), we leveraged DCC (15) and CIRCexplorer2 tools (14), which allow for fast and robust detection and quantification of circRNAs by identifying non-canonical back-splice junction reads with a low false-positive rate (33, 34). With a cut-off that back-splicing reads were greater than two in at least two samples, we identified 6726 and 5707 circRNAs by using DCC and CIRCexplorer2, respectively (**Figure 1b**). For the 4902 circRNAs detected by both algorithms, 99.8% of them could be annotated according to the circRNA databases, i.e., circBase (17), circAtlas v2 (35), and CIRCpedia v2 (36), and 11 novel circRNAs were unraveled in this study (**Figure 1c**).

### Molecular features of the circular RNAs in human skin wounds

We examined if there was any global change in circRNA production during wound repair or in chronic wounds. Comparing the four sample groups, i.e., the skin, Wound1, Wound7, and VU, we did not find obvious difference in regard to the number of exons occupied by circRNAs, the putative spliced length of circRNAs, or the number of circRNAs generated per gene (**Figure S1a-c**, **Dataset S1**). For example, we show that the circRNAs detected in VU samples occupied four exons on average and 37 exons at a maximum; their average spliced length was 681nt and the longest one was 7239nt; one to twelve (a mean value of 1.57) circRNA variants were derived from one gene, and most of genes (∼1400) only generate one circRNA variant. Moreover, we found that the total abundance of circRNAs was similar for the Skin, Wound7 and VUs, while there was a slight increase in the Wound1 compared to the skin (**Figure S1d**).

We further interrogated the interplay between canonical splicing and back-splicing for the circRNAs in human skin wounds. In line with the canonical splicing rules (37), we observed that circRNAs barely made use of the first exon of the corresponding linear transcript as starting circularized exon. Approximately 33% circRNAs detected in this study favored the second exon as a starting point (**Figure S1e**). However, for the ending circularized exon, we could not see such a clear preference (**Figure S1f**). Notably, almost all the circRNAs detected here (4900/4902) are exonic circRNAs (**Dataset S1**). Interestingly, we found that the flanking introns of circRNAs (upstream, downstream, or both sides) were significantly (Mann-Whitney-Wilcoxon Test, *P* < 0.001) longer than the introns genome-wide (**Figure 1d**). This may be due to that the longer flanking introns contain more repetitive complementary elements that facilitate circRNA biogenesis in mammalian cells (38, 39). By using rMATS tools, we showed that exon skipping (∼70.7%) was the most predominant among five main types of alternative splicing (AS) events (20) (**Figure S2a**). After correcting for intron lengths and gene expression levels, we found that AS events were significantly more common among the circRNA-generating exons than the non-circRNA-generating exons in all four sample groups (**Figure 1e**, **Figure S2b-d**). In addition, circRNA-forming exons showed higher exon inclusion levels compared to exons that do not form circRNAs, suggesting that circRNAs generation is not a byproduct of exon skipping (**Figure 1f**, **Figure S2e-g**).

We further profiled the circRNA expression in two subcellular compartments of human primary epidermal keratinocytes, i.e. nucleus and cytosol. We detected a total number of 3858 circRNAs expressed in keratinocytes out of which 1833 overlapped with the circRNAs identified in the human wound samples (**Figure 1g**). 3538 circRNAs were exclusively detected in the cytosol compartment of keratinocytes (**Figure 1h, Dataset S2**), which is in line with the current knowledge that exonic circRNAs are preponderantly found in the cytosol of cells (40).

Together, the circRNAs detected in each sample group displayed similar molecular features, thereby we postulated that the global production of circRNAs was not altered during wound healing or in VU.

### CircRNAs differentially expressed during wound repair and in VU

Our next goal was to identify individual circRNAs that were specifically regulated in the healing or non-healing human wounds. Based on the circRNA expression profiles, the four sample groups, i.e., the skin, Wound1, Wound7, and VU, were clearly separated in the principal component analysis (PCA) (**Figure 2a**). Especially, we identified 97 differentially expressed (DE) circRNAs by pairwise comparison [fold change (FC) ≥ 2, FDR < 0.05 in at least one of the six comparisons of any two groups] (**Figure 2b**, **Table S5**). Interestingly, only weak correlations (Pearson coefficient: 0.03 ∼ 0.08) were found between circRNA and their cognate linear RNA absolute expression read counts in each sample group, suggesting that circRNA expression was largely independent of their cognate linear RNA expression (14, 41) (**Figure S3**). Moreover, when comparing chronic or acute wounds with the skin, we found that the overall expression of circRNAs was upregulated (mean log_2_ FC= 0.03 – 0.27) while linear RNAs were downregulated (mean log_2_ FC= -0.13 – -0.09) (**Figure 2c**). For the 97 DE circRNAs, we observed that their cognate linear RNAs were either altered in the same direction as circRNAs or not significantly changed in the respective comparison (**Figure 2c, Table S5, Dataset S3**). Together, the different expression patterns of circRNAs compared to their cognate linear RNAs suggested that the circRNA expression was subjected to specific regulation, and therefore they may play functional roles in wound repair.

**Figure 2.**
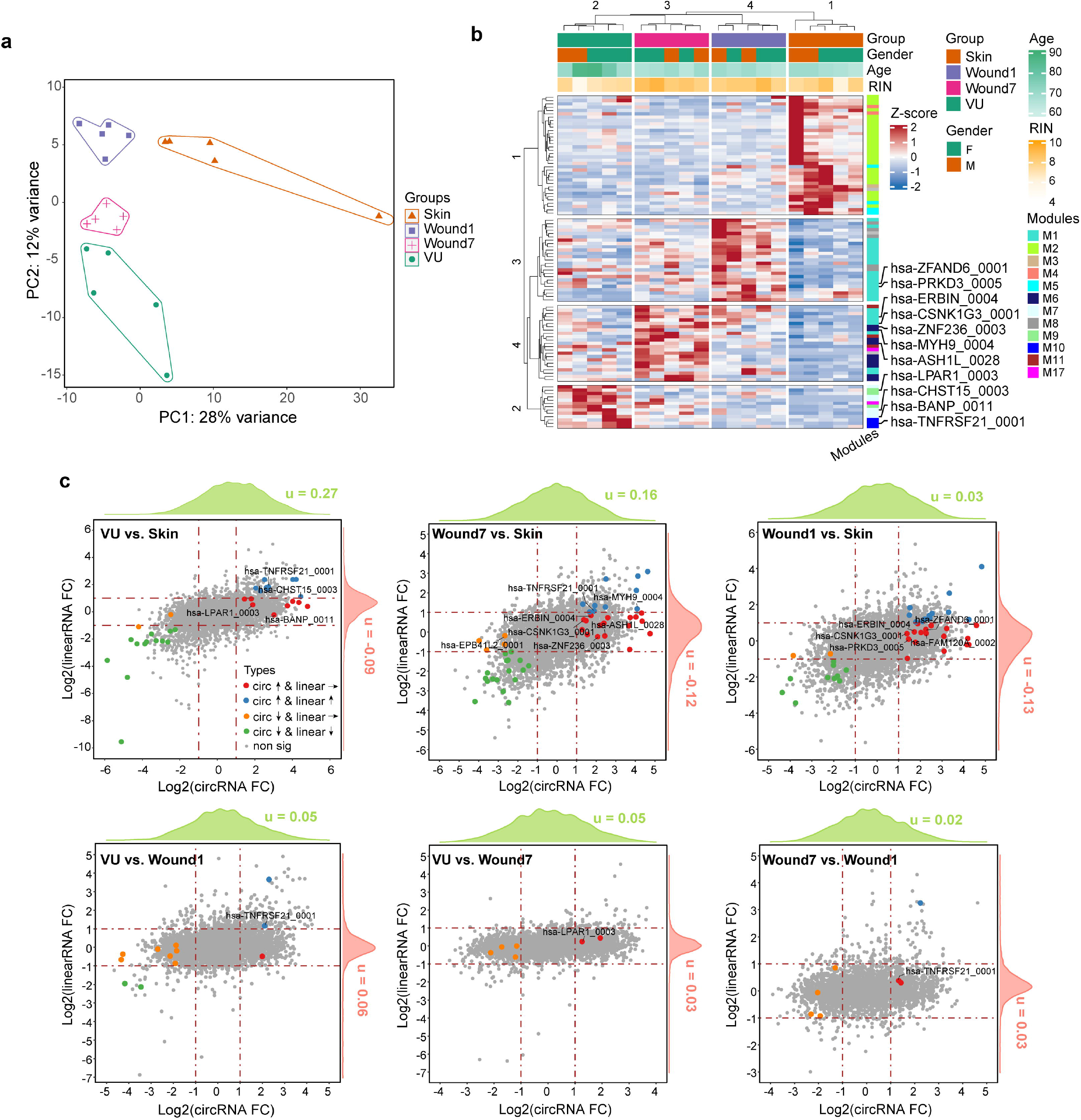
Differentially expressed circRNAs in human skin wounds. (**a**) PCA plot based on circRNA expression profile. Each dot represents an individual sample. (**b**) K-means clustering of the 20 sequenced samples based on 97 differentially expressed circRNAs. Donor’s sex, age, RNA quality (RIN) of each sample, and WGCNA modules (M) are depicted with color bars. (**c**) Pairwise comparison of circRNAs and their cognate linearRNAs’ expression. Differentially expressed circRNAs (FC ≥ 2, FDR < 0.05) are highlighted with colored dots according to the expression change patterns of their cognate linearRNAs. The experimentally validated circRNAs are labeled in the plots.

### Co-expression networks of circRNAs and mRNAs in human skin wounds

Besides the DE analysis, we performed weighted gene co-expression network analysis (WGCNA) using the RNA-seq data of both circRNAs and mRNAs. This approach allowed us to identify circRNAs and mRNAs potentially important for each phenotypic trait, in this case, the Skin, Wound1, Wound7, or VU (24, 42). Also, based on a hypothesis that the co-expressed genes may be involved in the same or related biological processes (43), one may catch a first glimpse on the putative function of circRNAs by looking at the co-expressed mRNAs. We identified 18 distinct modules (M) summarized by the module eigengenes (MEs) that represents the first principal component of both the circRNA and mRNA expression in each module (**Figure 3a, b**, **Dataset S4**). The robustness of these modules was confirmed by module preservation analysis (**Figure S4b**). Next, we performed gene ontology (GO) analysis for the mRNAs in the modules that were significantly (Pearson’s correlation, FDR < 0.05) correlated with each sample group (**Figure 3c**, **Figure S4c**). With this approach, the circRNAs that were closely related to the fundamental biological processes essential for wound healing were unraveled (**Dataset S4**). We found that the intact skin associated circRNA and mRNA co-expression networks (M3, M2, and M5), which were mainly required for the skin barrier functions, were downregulated in both acute and chronic wounds. In the inflammatory phase acute wounds (Wound1), we identified 474 and 75 circRNAs’ expression correlated with the mRNAs in M1 and M8 modules, respectively, which were involved in innate immune response. Moreover, in the proliferative wounds (Wound7), 123 circRNAs were found co-expressed with the M6 mRNAs important for cell cycle. Importantly, we added a list of circRNAs to the pathological molecular signature of VU, including increased extracellular matrix organization and cell adhesion and abnormal chronic inflammation (M7 and M10). Overall, the co-expression network modeling helped to sift circRNAs with putative functions underlying healing and non-healing human wounds, which lays a step stone for further functional study of circRNAs in wound repair.

**Figure 3.**
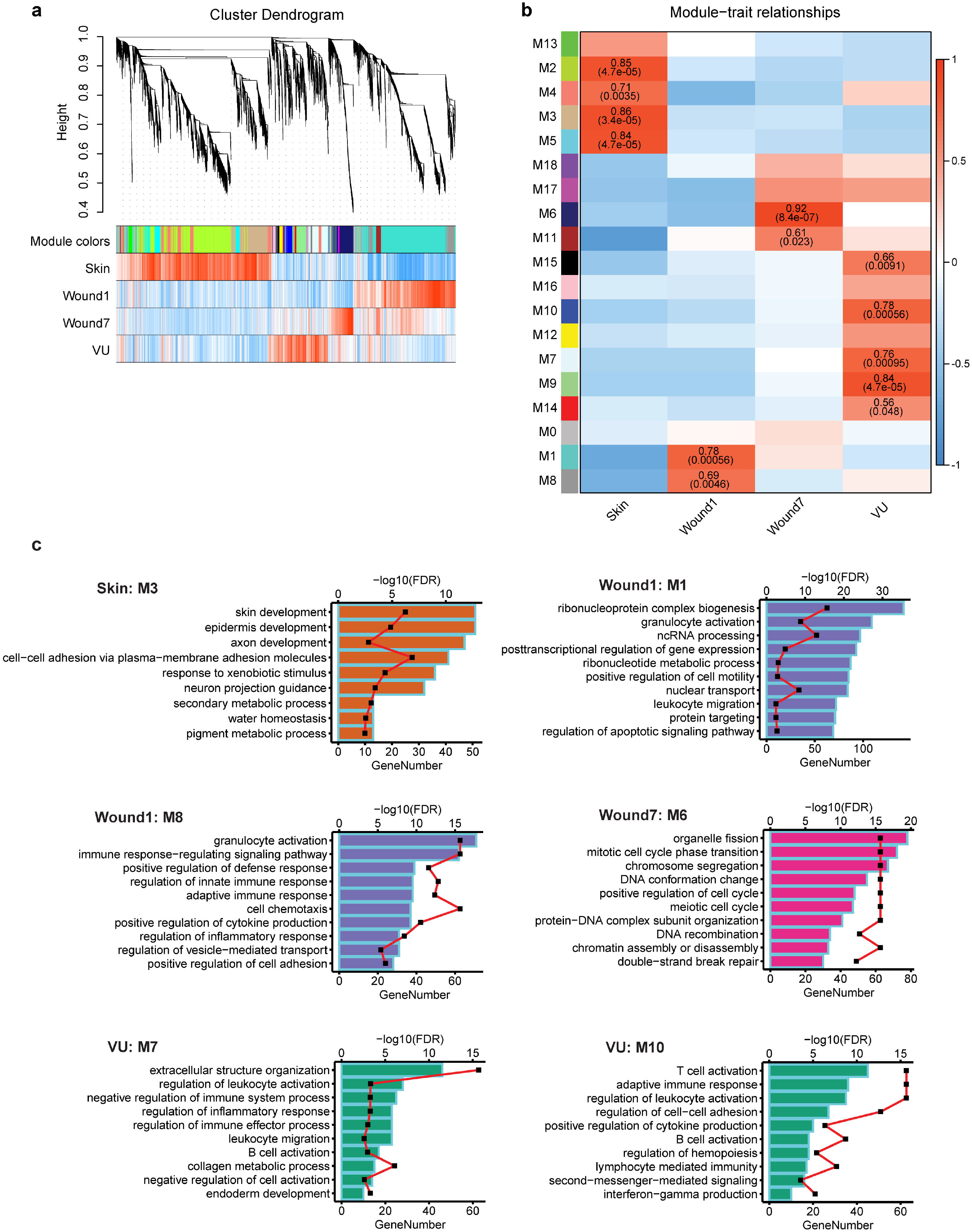
Weighted gene co-expression network analysis (WGCNA) of circRNAs and mRNAs in human skin wounds. (**a**) Cluster dendrogram of co-expression modules based on the combined circRNA and mRNA expression profiles. Each branch corresponds to a module colored in bar plots (the 1^st^ row) and each leaf indicates a circRNA or mRNA. Pearson correlation coefficients between each sample group and RNA are shown in bar plots in the 2^nd^ to the 5^th^ row (red and blue lines represent positive and negative correlations, respectively). (**b**) Pearson correlations between distinct module eigengenes (ME) and traits (sample groups) are depicted in a heatmap. The correlation coefficients and (FDR values) are labeled in the significant modules (FDR < 0.05). (**c**) Top ten enriched gene ontology (GO) terms ranked by Gene number in each GO (bar length, bottom x axis) with FDR less than 0.01 (red line, top x axis) for the significant modules are shown in bar plots.

### CircRNAs serving as miRNA sponge in wound healing

Many circRNAs have been shown to act as miRNA sponges, restraining miRNAs from repressing their target mRNAs (44–46). To identify the putative miRNA-sponging circRNAs in the wound healing context, we constructed circRNA-miRNA-mRNA networks by integrating the current circRNA and mRNA expression profiles with the miRNA sequencing data of the same samples. First, we searched for miRNA target sites in both circRNAs and mRNA sequences. Second, we sifted the connections between the circRNA, miRNA, and mRNA nodes based on the criteria of significantly negative correlations of circRNA-miRNA and miRNA-mRNA interactions, and positive correlations of circRNA-mRNA interactions (Spearman’s P value < 0.05) (**Figure 4a**). Among the co-expression network modules significantly associated with the Skin, Wound1, Wound7, and VU (**Figure 3b, Dataset S4**), we shortlisted the positively corelated circRNAs and mRNAs that could compete for common miRNAs (**Figure 4b**, **Figure S5-7**). For example, hsa-CHST15_0003 may enhance the expression of SH2B3 through sponging miR-125b-5p and let-7a (**Figure 4b**). SH2B3 has been previously identified as a negative regulator of wound healing by changing the extracellular matrix (ECM) production, angiogenesis, and inflammation (47, 48). Here we found that both hsa-CHST15_0003 and SH2B3 expression were upregulated in VU, while miR-125b-5p and let-7a levels were lower in VUs. The circRNA-miRNA-mRNA network analysis identified the circRNAs that may act as miRNA sponge, which could explain some circRNAs and VU pathology-relevant mRNAs’ co-expression patterns.

**Figure 4.**
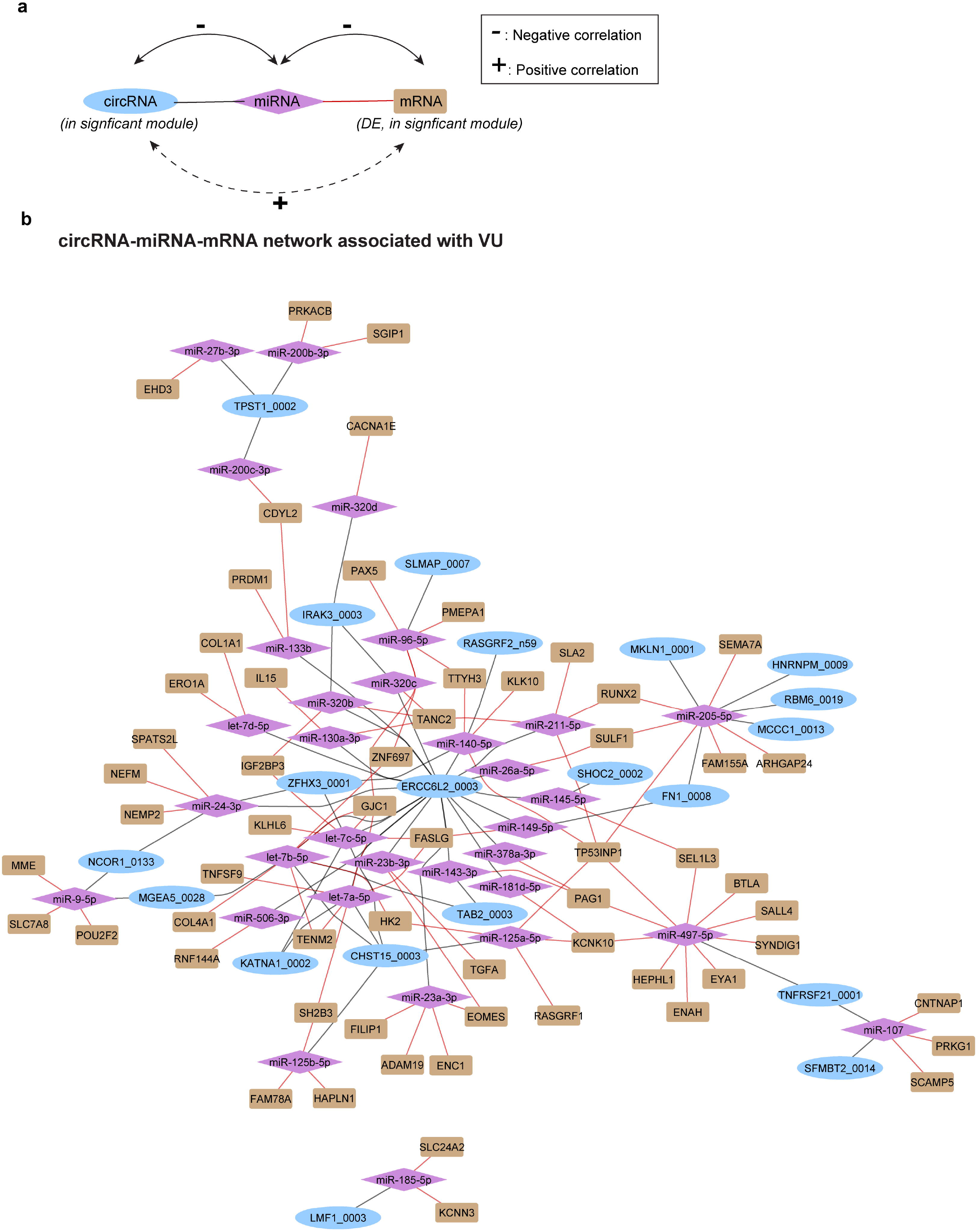
CircRNA-miRNA-mRNA networks in human skin wounds. (**a**) Schematic diagram showing the criteria of constructing circRNA-miRNA-mRNA interactions. “-” and “+” represent negative and positive correlations of their expression profiles by using Spearman’s correlation tests. (**b**) Networks linking the circRNAs and mRNAs co-expressed in VU-associated modules via their interactions with miRNAs.

### Experimental validation of the *in silico* identified circRNAs in human wounds

We randomly selected 11 circRNAs from the 97 DE circRNAs (**Figure 2b**) and two circRNAs without significant expression change (i.e., hsa-TNPO2_0001 and hsa-VPS13D_0016) to validate their circularity (**Figure 5b, Figure S8**). For this, we performed RT-PCR with outward-facing primers in total RNA isolated from human epidermal keratinocytes and dermal fibroblasts (**Figure 5a**, **Table S6**), and then Sanger sequencing of the RT-PCR products. We confirmed the back-splice junction sites (BSJ) of these 13 circRNAs identified in the *in silico* analysis of the RNA-seq data (**Figure 5b, c**). Moreover, we treated the RNA samples with RNase R, which efficiently digested linear RNAs, e.g., 18S and 28S ribosomal RNA (**Figure 5d**). RT-PCR analysis showed that hsa-TNFRSF21_0001, hsa-CHST15_0003, hsa_ASH1L_0028 and hsa_BANP_0011 were completely resistant to the degradation by RNase R, further confirming their circularity (**Figure 5e**) (40). Of note, hsa_ZFAND6_0001, hsa_ZNF236_0003, hsa_ERBIN_0004, hsa_CSNK1G3_0003, and hsa_TNPO2_0001 were partially degraded by RNase R, which was in line with previous studies showing that some circRNAs are less RNase R-resistant (49). We further validated six DE circRNAs using quantitative RT-PCR on an extended cohort including matched skin and acute wounds from five healthy donors and 12 VUs, which reproduced their expression patterns from RNA-seq (**Figure 5f**). Together, these experimental results support the robustness and reproducibility of our in-silico analysis of RNA-seq data.

**Figure 5.**
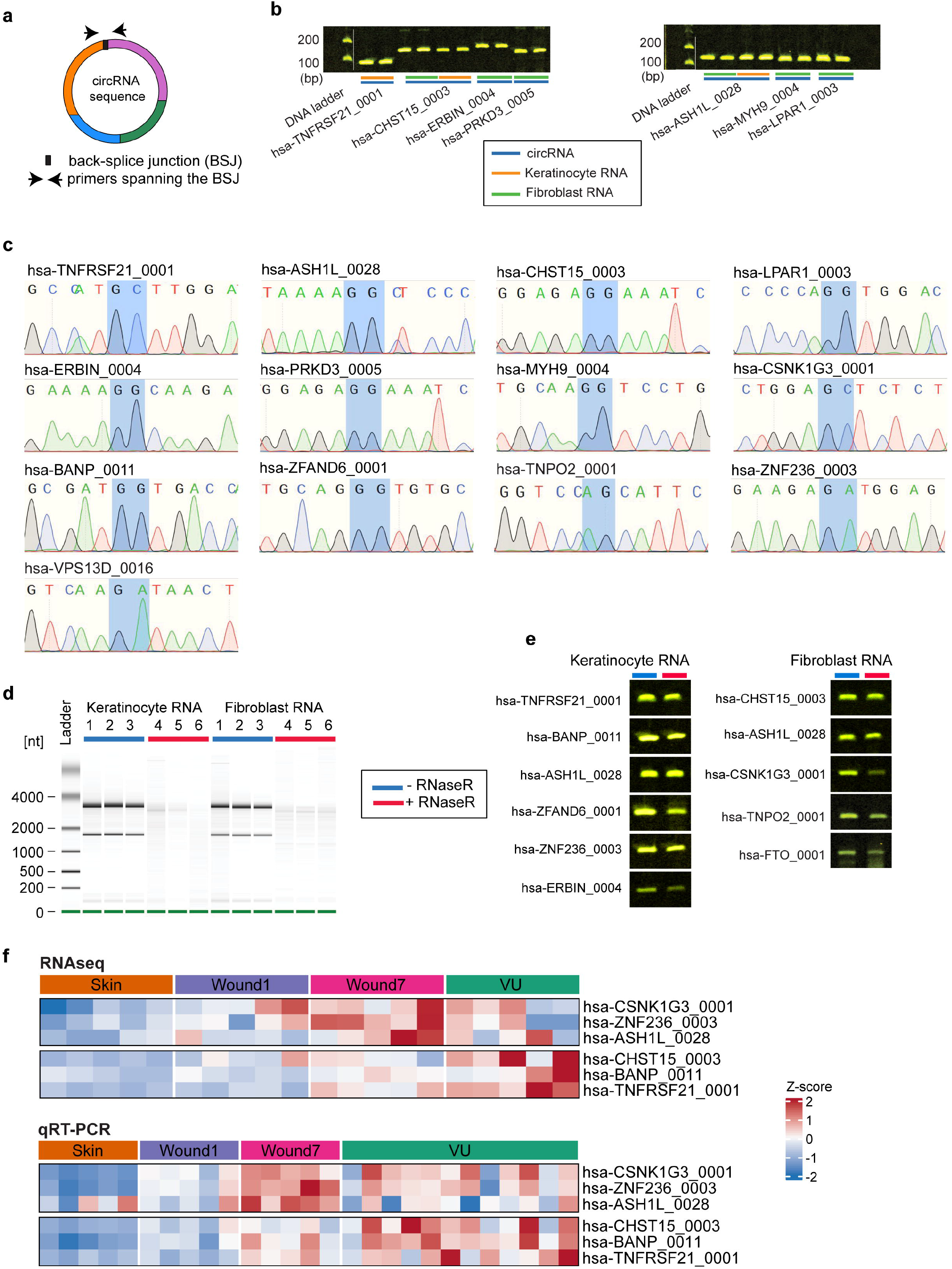
Experimental validation of the in silico identified circRNAs in human wounds. (**a**) Illustration of primer design to amplify the BSJ region of a circRNA. (**b**) Agarose gel electrophoresis of the PCR products on cDNA amplified with specific divergent primers for the candidate circRNAs. (**c**) Sanger sequencing confirming the BSJ sequence (shaded in blue) for the candidate circRNAs. (**d**) Simulated gel image generated by a Bioanalyzer showing the absence of the ribosomal RNA 18S and 28S bands after RNaseR digestion. (**e**) Agarose gel electrophoresis of the PCR products on cDNA obtained from RNaseR-digested RNA (red) or control RNA (blue) from keratinocytes or fibroblasts. (**f**) Heatmaps showing the circRNAs expression patterns from RNAseq and the validation by qRT-PCR in an extended cohort of paired Skin, Wound1, and Wound7 samples (n=5) and VUs (n=12). 18S was used for normalization. Z scores show the normalized circRNA expression across different sample groups.

### Hsa-TNFRSF21_0001 and hsa-CHST15_0003 regulate keratinocyte functions important for wound healing

Next, we asked whether the circRNAs with changed expression may play a functional role in wound healing. As a proof-of-concept, we dug into the biological functions of two circRNAs i.e., hsa-TNFRSF21_0001 or hsa-CHST15_0003, which were expressed in keratinocytes and upregulated in VU (**Figure 2b**, **Figure 5b**). We knocked-down their expression in human epidermal keratinocytes by transfecting BSJ-specific siRNAs (**Figure S9**) and then performed transcriptomic analysis by microarray. Gene set enrichment analysis (GSEA) of the transcriptomic changes upon the depletion of hsa-TNFRSF21_0001 or hsa-CHST15_0003 helped us gain a first insight into the functions of these two circRNAs (**Dataset S5**). We found that knockdown of hsa-TNFRSF21_0001 upregulated the genes related to epidermal cell differentiation, epithelial-to-mesenchymal transitions (EMT) and adhesion, while downregulating the genes important for cell cycle (**Figure 6a**). The expression changes and interactions of the genes involved in these cellular functions were denoted in the functional networks (**Figure 6b-d**). We further confirmed the hsa-TNFRSF21_0001-mediated regulation of several network hub genes related to keratinocyte EMT and cell adhesion *(ACTA2*, *ITGB1,* and *VEGFA)*, differentiation (*IVL*), and cell cycle (CCNE1) by qRT-PCR (**Figure 6e**). GSEA showed that silencing hsa-CHST15_0003 expression also led to increased expression of cell migration related genes but lower levels of cell cycle related genes (**Figure 7a-c**). By qRT-PCR, we validated that expression of the hub genes in the functional networks underlying cell proliferation (TIMP1 and VEGF) and migration (E2F1 and CCNE1) were regulated in the opposite directions by hsa-CHST15_0003 (**Figure 7d**). In line with the gene expression changes, by live-cell imaging, we found that silencing either hsa-TNFRSF21_0001 or hsa-CHST15_0003 promoted keratinocyte migration whereas delaying their growth (**Figure 7e-h, Suppl. Video**).

**Figure 6.**
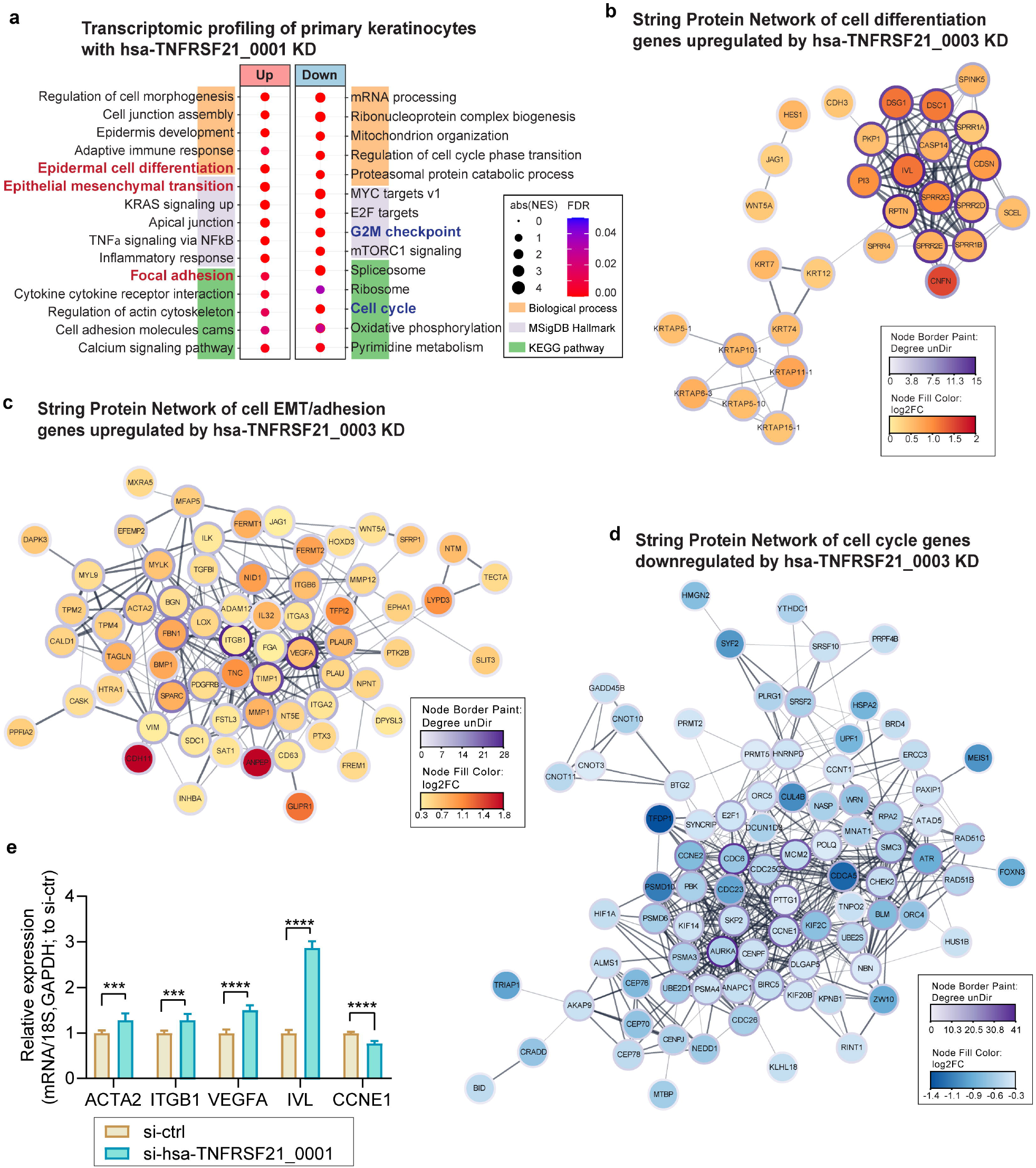
Transcriptomic analysis of keratinocytes with hsa-TNFRSF21_0001 knockdown. (**a**) Transcriptome profiling in keratinocytes 24h after hsa-TNFRSF21_0001 knockdown (KD). Gene Set Enrichment Analysis (GSEA) of the upregulated or downregulated genes by hsa-TNFRSF21_0001 KD in terms of biological processes, MSigDB hallmarks, and KEGG pathways (**b-d**). Functional association networks are presented for the genes related to (**b**) Epidermal cell differentiation, or (**c**) Epithelial mesenchymal transition and focal adhesion KEGG pathway, or (**d**) G2M checkpoint and Cell cycle KEGG pathway, highlighted in (**a**). Core nodes are identified by centrality analysis and marked as indicated. (**e**) qRT-PCR validation of hub gene expression after hsa_TNFRSF21_0001 KD. *18S* and *GAPDH* mRNA levels were used for normalization (n=4). The data for each sample group is shown as relative to si-ctrl sample levels and presented as mean ± SD. *** P < 0.001 by unpaired two-tailed Student’s t test.

**Figure 7.**
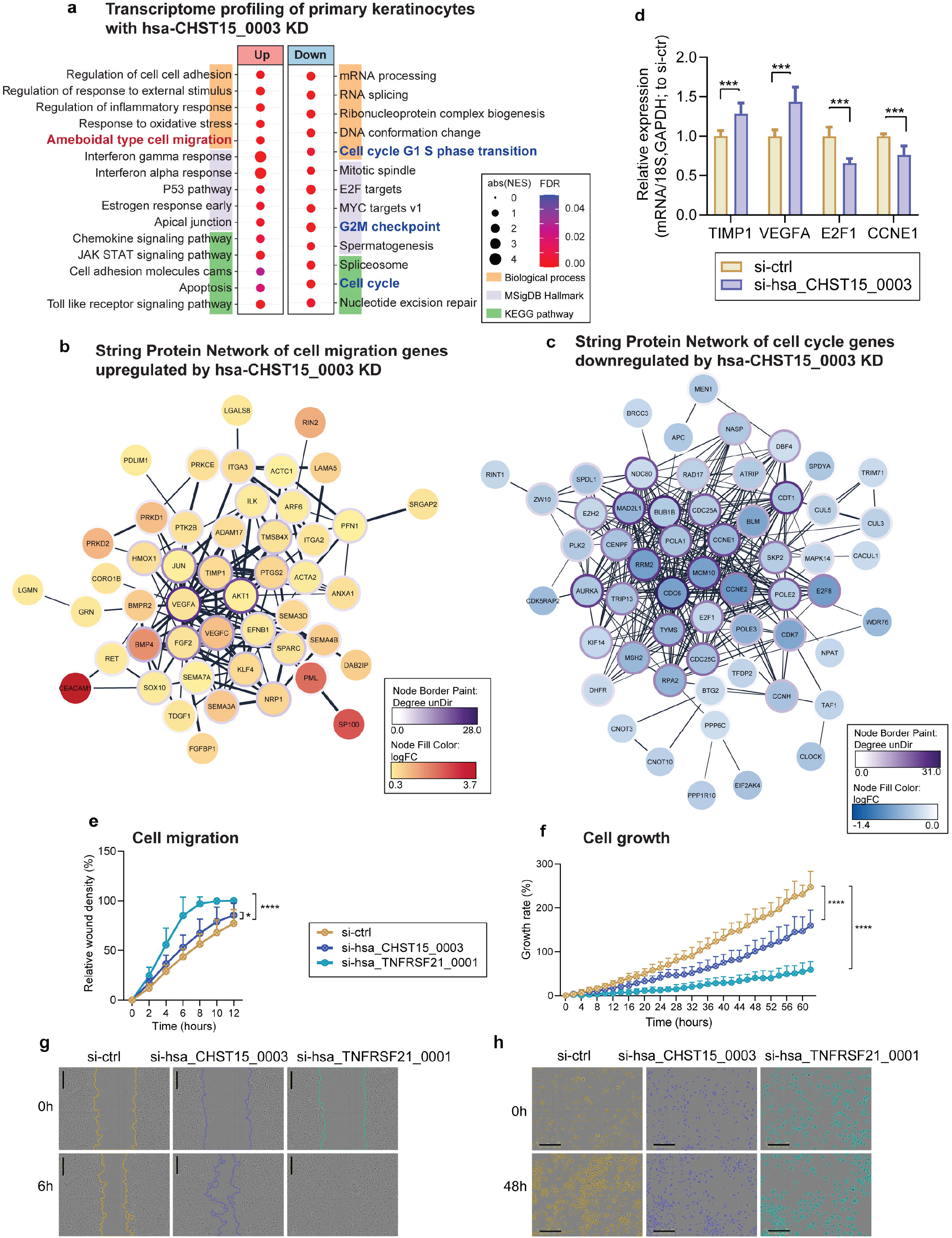
hsa-TNFRSF21_0001 and hsa-CHST15_0001 regulate keratinocyte functions important for wound healing. (**a**) Transcriptome profiling in keratinocytes 24h after hsa-CHST15_0003 knockdown (KD). Gene Set Enrichment Analysis (GSEA) of the upregulated or downregulated genes by hsa-CHST15_0001 KD in terms of biological processes, MSigDB hallmarks, and KEGG pathways (**b, c**). Functional association networks are presented for the genes related to (**b**) Ameboidal cell migration, or (**c**) Cell cycle G1-S phase transition, G2M checkpoint and Cell cycle KEGG pathway, highlighted in (**a**). Core nodes are identified by centrality analysis and marked as indicated. (**d**) qRT-PCR validation of hub gene expression after hsa_CHST15_0003 KD. *18S* and *GAPDH* mRNA levels were used for normalization (n=4). The data for each sample group is shown as relative to si-ctrl sample levels. (**e**) Live-cell imaging using the IncuCyte^®^ system of migrating (**e, g**) or proliferating (**f, h**) keratinocytes with either hsa-TNFRSF21_0001 or hsa-CHST15_0003 KD (*n*L=L10). (**e**) Plot shows the cell migration rates. (**f**) Plot shows cell proliferation rates based on cellular confluence. Representative photographs of migrating (**g**) or proliferating (**h**) keratinocytes at the indicated timepoints. Scale barL= 300 μm. Data are presented as mean ± SD (**d, e, f**). *** P < 0.001 by unpaired two-tailed Student’s t test (**d**), * P< 0.05, **** P < 0.0001 by two-way RM ANOVA + Dunnett’s test (**e, f**).

## Discussion

Here we performed a comprehensive evaluation of the genome-wide expression of circRNAs during human skin wound healing and in chronic non-healing VU. Although changed overall circRNA production has been reported in epidermal differentiation (7), psoriasis, atopic dermatitis (50), and squamous cell carcinoma (51), our data showed that circRNAs were not globally altered during human skin wound healing or in VU. Instead, we found that the circRNA expression patterns were dynamically changed with the healing process, and the chronic wound exhibited an abnormal circRNA signature. Interestingly, different expression patterns were found between circRNAs and their cognate linear RNAs, suggesting that the circRNA production is subjected to specific regulation during wound repair and they may play functional roles in this process.

To decode the putative physiological and pathological roles of circRNAs, we performed co-expression network modeling to identify the key circRNA and mRNA modules underpinning each healing stage and chronic wound pathology. Integration of circRNA and protein-coding gene expression indicated the biological processes that circRNAs may be involved in. Moreover, incorporation of the paired miRNA expression profiles into these circRNA-mRNA networks exposed the circRNAs that could sponge miRNAs and rescue the expression of the miRNA target genes (6). Some co-expressed circRNAs and mRNAs may be linked via regulation by miRNAs. While the competing endogenous RNA mechanism only partly reflects the diverse mechanistic spectrum of circRNAs, other modes of action, such as interaction with proteins, cannot be inferred from the current transcriptomic data. Nevertheless, the circRNA-miRNA-mRNA networks specific to different healing stages or non-healing states of human wounds *in vivo* enables us to sift clinically relevant circRNAs for further in-depth functional and mechanistic studies.

Importantly, our study unravels the pathological roles of hsa-TNFRSF21_0001 and hsa-CHST15_0003 in human VU. Both circRNAs could repress the migration of epidermal keratinocytes while increasing their proliferation. Also, hsa-TNFRSF21_0001 depletion upregulated many differentiation-related genes in keratinocytes. Thereby, increased expression of these two circRNAs in VUs may be partially responsible for the nonmigratory, hyperproliferative, and abnormally differentiated state of the VU wound-edge keratinocytes (52). Diving into the functional networks consisting of the genes regulated by these circRNAs, we identified several hub genes that may possess a greater regulatory ability over the other genes in the network (53) and explain the observed biological roles of these circRNAs. For example, we showed that silencing hsa-TNFRSF21_0001 enhanced the expression of the keratinocyte differentiation marker *IVL, as well as ITGB1* that is essential for basement membrane formation, cell adhesion, and migration (54, 55).

On the contrary, silencing either of these two circRNAs decreased the expression of CCNE1 – a master regulator of cell cycle progression that was previously identified as a key player in keratinocyte hyperproliferation in psoriasis (56, 57). Similarly, upon hsa-CHST15_0003 silencing, the expression of *E2F1* was decreased, which gene is important for maintaining epidermal keratinocytes in a proliferative but undifferentiated state (58). We also found that hsa-CHST15_0003 negatively regulated *TIMP1* expression. As one member of the tissue inhibitors of metalloproteases, *TIMP1* modulates the breakdown of the extracellular matrix and cytokines of the wound environment, thus affecting cell migration in wound healing (59). Abnormal *TIMP1* expression has been associated with delayed healing, which may be partially explained via hsa-CHST15_0003 dysregulation in keratinocytes at non-healing wound-edge.

Our study however does not account for the cellular heterogeneity of human wounds, and the possibility that the altered circRNA expression is partially due to changes in cell-type proportions during wound healing cannot be excluded. This limitation could be addressed by detecting circRNA expression of single cells, such as by in situ hybridization of individual circRNA or single-cell total RNA sequencing (sc total RNA-seq) with high sensitivity to non-poly(A) RNA. The sc total RNA-seq technology is emerging but is still suffering from setbacks such as insufficient sequencing depth and lack of bioinformatic tools, it is thus not ready to be used for single-cell circRNA exploration in complex tissues (60) (61).

## Conclusions

In conclusion, our study unraveled the dynamically changed circRNA-miRNAs-mRNA networks underpinning each phase of human skin wound healing and VU pathology. We shortlisted the most clinically relevant circRNAs for further studies on their function and mechanism in wound repair. In addition, we unraveled the pathological roles of hsa-CHST15_0003 and hsa-TNFRSF21_0001 in chronic non-healing VU. Our work paves the way to decipher functional significance of circRNAs in tissue repair.

## List of abbreviations

AS: Alternative splicing
BH: Benjamini-Hochberg
BP: Biological process
circRNA: circular RNA
DE: Differentially expressed or differential expression
ECM: Extracellular matrix
EMT: Epithelial-to-mesenchymal transitions
FDR: False discovery rate
FPKM: Fragments per kilobase of transcript, per million mapped reads
FPM: Fragments mapped to back-splicing junctions, per million mapped fragments
GEO: Gene Expression Omnibus
GO: Gene ontology
GSEA: Gene Set Enrichment Analysis
MEs: Module eigengenes
miRNA: microRNA
mRNA: messenger RNA
PCA: Principal component analysis
siRNA: small interfering RNA
VU: Venous ulcer
WGCNA: Weighted gene co-expression network analysis

## Availability of data and materials

The data of the RNA sequencing and microarrays performed in the current study are available in the NCBI Gene Expression Omnibus (GEO) database under the accession number GSE174661 and GSE188939, respectively. In addition, the analyzed dataset is presented with an online R Shiny app and can be accessed through a browsable web portal (http://130.229.28.87/shiny/circRNA_wholebiopsy-shinyApp/). Complementary data supporting the conclusions of our study are presented in additional files: Dataset S1 – Identification of circRNAs in human skin wounds; Dataset S2 – Subcellular localization of circRNAs in keratinocytes; Dataset S3 - Pairwise comparison of mRNA expression in human skin wounds; Dataset S4 – WGCNA and GO analysis of circRNAs and mRNAs in human skin wounds; and Dataset S5 – GSEA of keratinocytes with hsa-TNFRSF21_0001 or hsa-CHST15_0003 knockdown.

## Code availability

The R code for the DE analysis and the WGCNA analysis is provided in Additional File: Code S1 and S2.

## Acknowledgements

We express our gratitude to the patients and healthy donors participating in this study. We thank Mona Ståhle, Desiree Wiegleb Edström, Peter Berg, Fredrik Correa, Martin Gumulka, Mahsa Tayefi for clinical sample collection; Helena Griehsel for technical support. We thank the Microarray core facility at Novum, BEA, which is supported by the board of research at KI and the research committee at the Karolinska hospital. The computations/data handling was enabled by resources in projects of sens2020010 and SNIC2019/8-262 provided by the Swedish National Infrastructure for Computing (SNIC) at UPPMAX, partially funded by the Swedish Research Council through grant agreement no. 2018-05973.

## Funding

This work was supported by Swedish Research Council (Vetenskapsradet, 2016-02051 and 2020-01400), Ragnar Söderbergs Foundation (M31/15), Welander and Finsens Foundation (Hudfonden), LEO foundation, Ming Wai Lau Centre for Reparative Medicine, and Karolinska Institutet.

## Authors’ information

### Contributions

NXL and ZL conceived and designed the study. PS collected most clinical samples with assistance of MAT and DL. MAT, QW, LZ and DL performed the experiments. ZL and MAT carried out bioinformatic analysis. MAT, ZL, and NXL wrote the manuscript, which was revised and approved by all authors.

### Corresponding authors

Correspondence to Ning Xu Landén or Zhuang Liu

## Ethics declarations

### Ethics approval and consent to participate

Written informed consent was obtained from all donors for the collection and use of clinical samples. The study was approved by the Stockholm Regional Ethics Committee. The study was conducted according to the Declaration of Helsinki’s principles.

### Consent for publication

Not applicable.

### Competing interests

There are no competing interests to declare.

## Notes

http://130.229.28.87/shiny/circRNA_wholebiopsy-shinyApp/

### Competing Interest Statement

The authors have declared no competing interest.

