## Supplemental figures and tables for "Circular RNA signatures of human healing and non-healing wounds"

Additional Material for:

This PDF file includes:

Figures S1-S9 (pages 2-9)
Tables S1-S6 (pages 10-15)

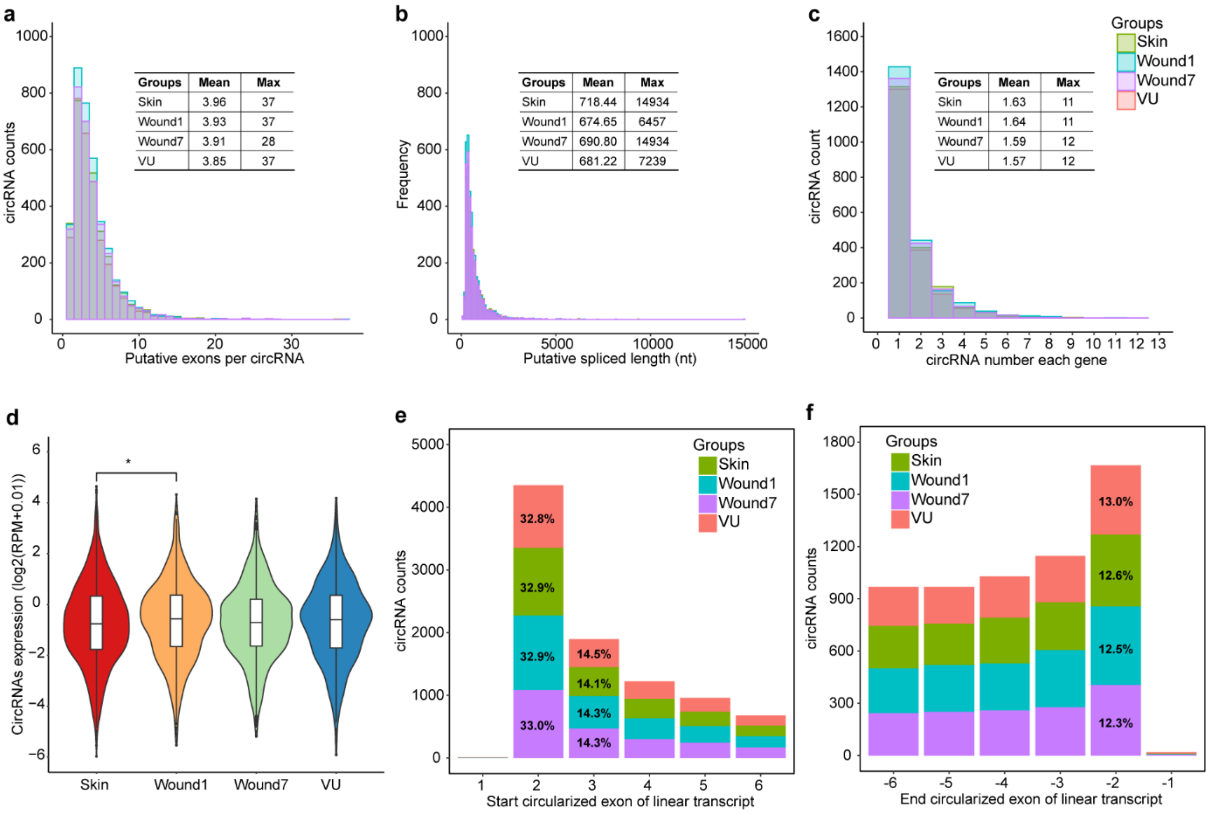


**Figure S1. Characterization of circRNAs detected in human wounds.** The distributions of putative exon numbers (**a**) and spliced lengths (**b**) of circRNAs in four groups (Skin, Wound1, Wound7 and VU). (**c**) The number of circRNAs generated per gene. Mean and maximum values of each group are shown in the plots. (**d**) Expressions of circRNAs in four groups. (**e, f**) Distributions of the first (**e**) and the last (**f**) six exons used for circRNA circularization. Mann-Whitney-Wilcoxon Test, **P* < 0.05 (**d**).

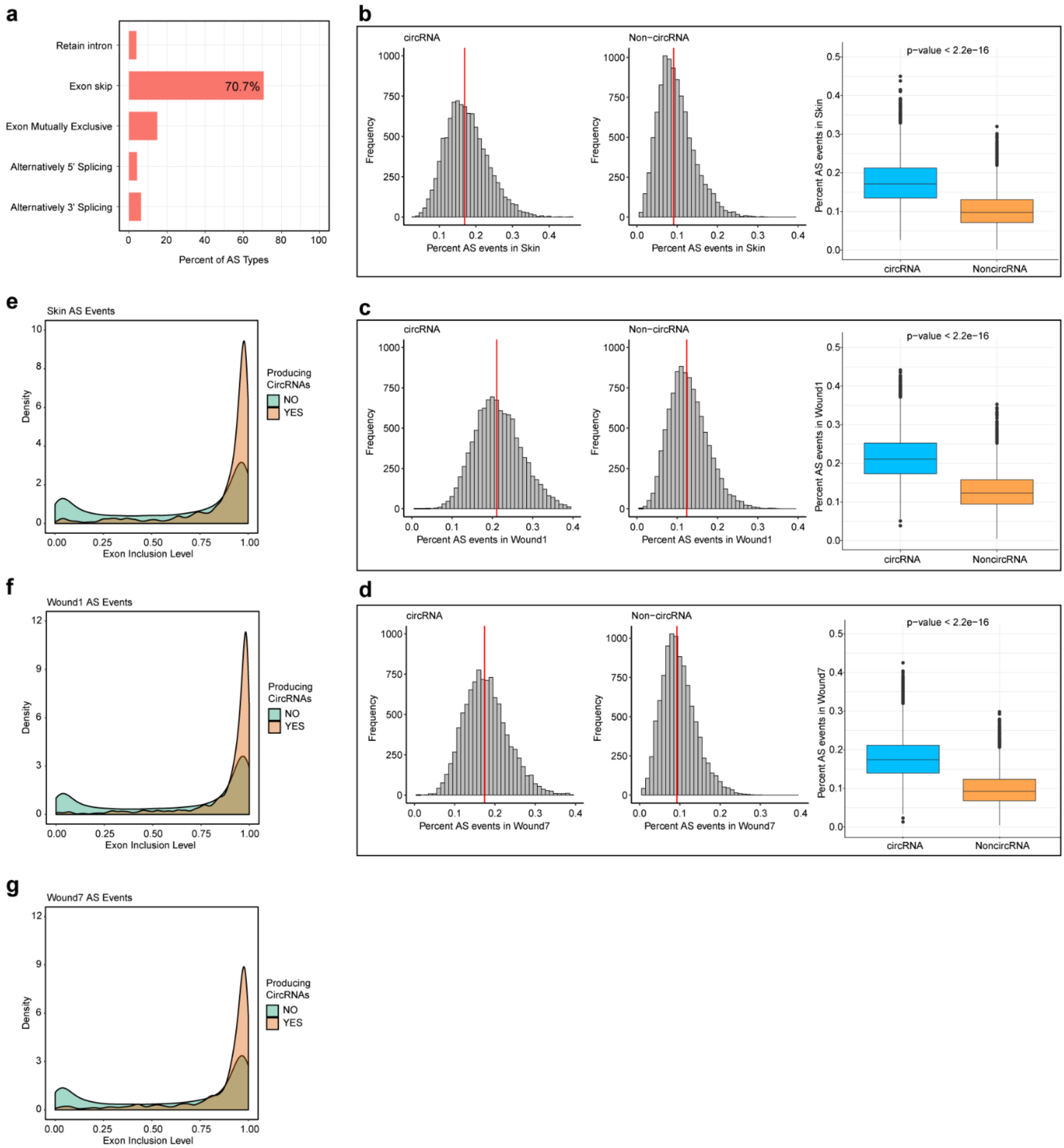


**Figure S2. Interplay between canonical splicing and back-splicing for circRNAs in human wounds.** (**a**) The average percentage of AS events of five main types detected in four groups. (**b, c, d**) Histograms showing the percentage of AS events detected in circRNA-forming exons (left) are significantly higher than in non-circRNA-forming exons (right) after 10000 random samplings corrected for intron lengths and gene expression levels in Skin (**b**), Wound1 (**c**), and Wound7 (**d**). (**e, f, g**) Density plots showing exon inclusion level for circRNA-forming exons are higher than non-circRNA-forming exons in Skin (**e**), Wound1 (**f**), and Wound7 (**g**). *P* values were calculated using Mann-Whitney-Wilcoxon Test.

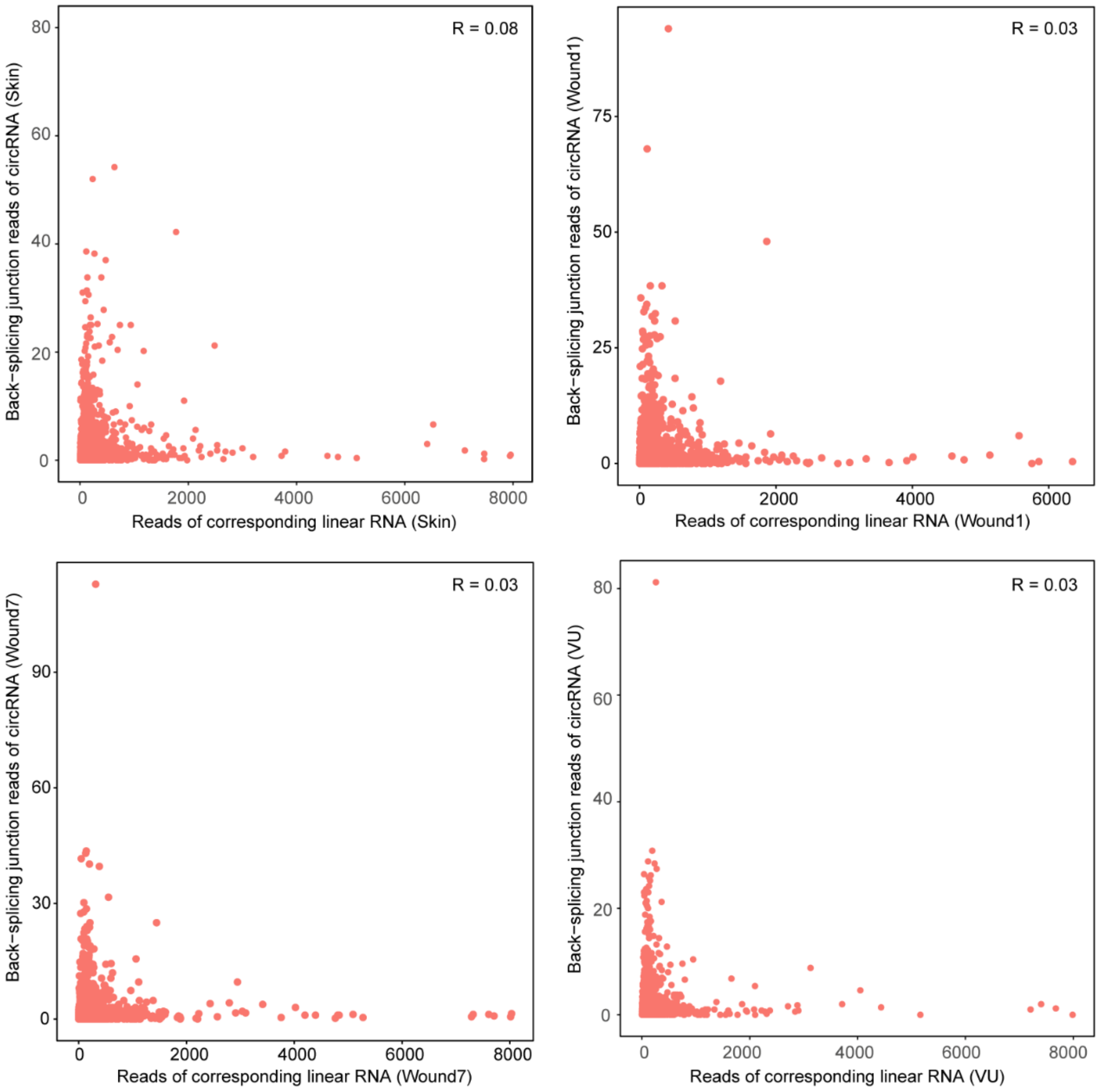


**Figure S3. The relationship between circRNA and their cognate linear RNA in Skin, Wound1, Wound7 and VU.** Dot plot of the Pearson correlation coefficients (R) between the absolute reads of circRNAs spanning the back-spliced junction sites and corresponding linear RNAs in four sample groups.

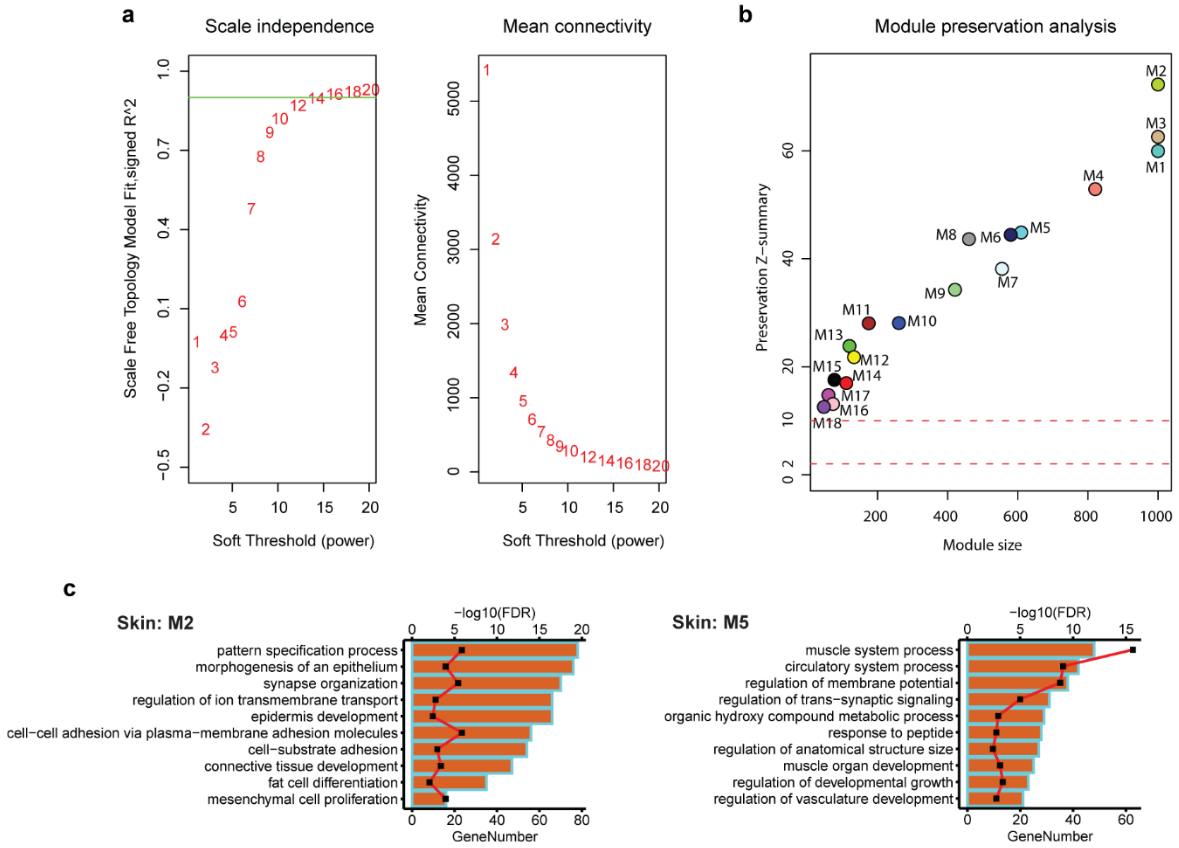


**Figure S4. Weighted gene co-expression network analysis (WGCNA) of combined circRNAs and mRNAs in human healing.** (**a**) Analysis of scale-free topology fit index (left panel) and mean connectivity (right panel) for selecting soft-threshold powers. Red line represents signed $R^2 = 0.8$. (**b**) Preservation analysis of co-expressed modules by permutating 200 times using the same 20 samples. Modular preservation is strong if Z-summary score > 10, weak to moderate if 2 < Z-summary score < 10, no evidence of preservation if Z-summary score ≤ 2. (**c**) Top ten enriched gene ontology (GO) terms ranked by Gene number in each GO (bar length, bottom x axis) with FDR less than 0.01 (red line, top x axis) for significant modules are shown in bar plots.

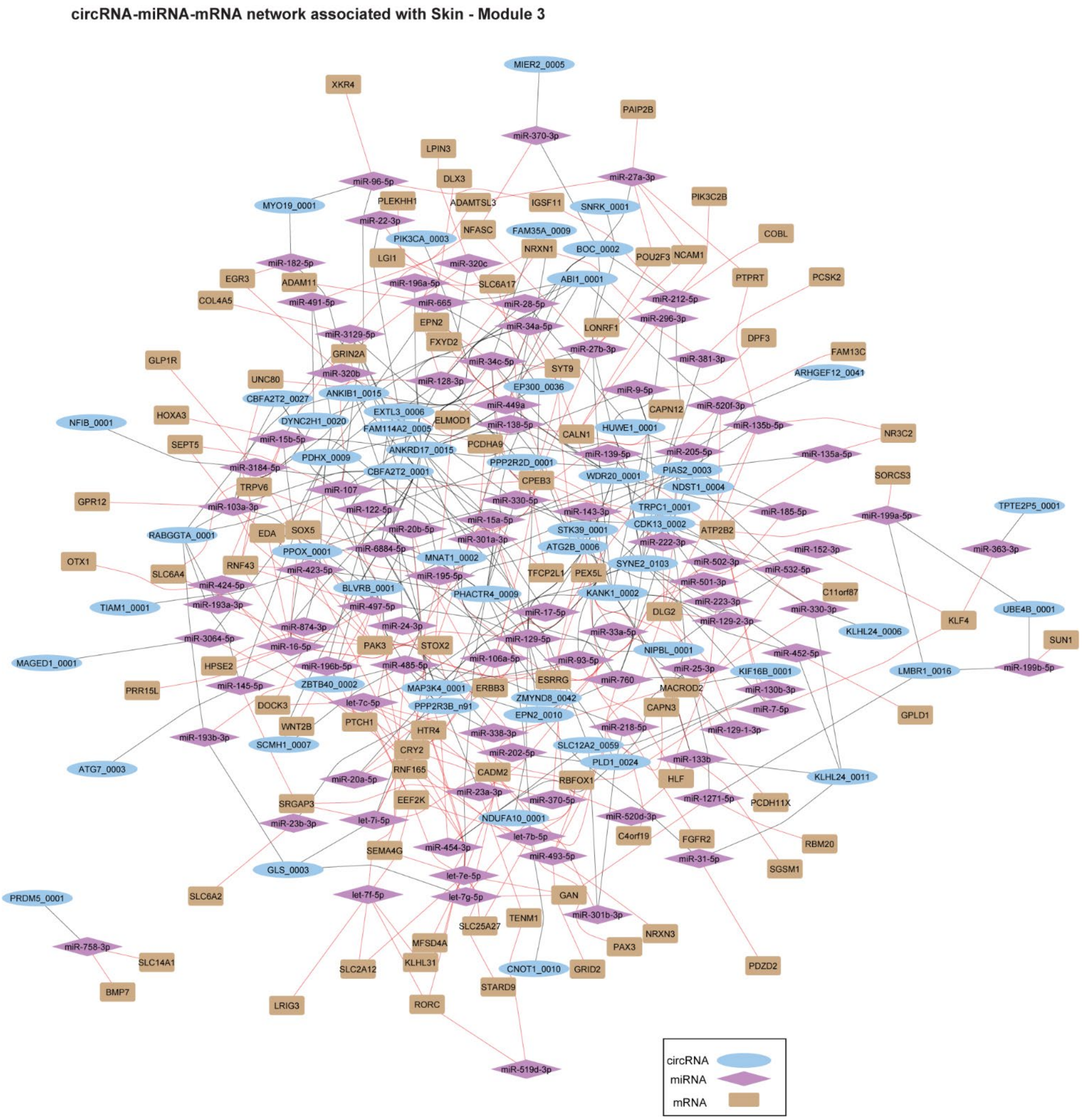


**Figure S5. CircRNA-miRNA-mRNA interactions in Skin-associated modules.** Interaction network of the circRNAs and mRNAs that are co-expressed in the Skin-associated module M3 and linked via miRNAs.

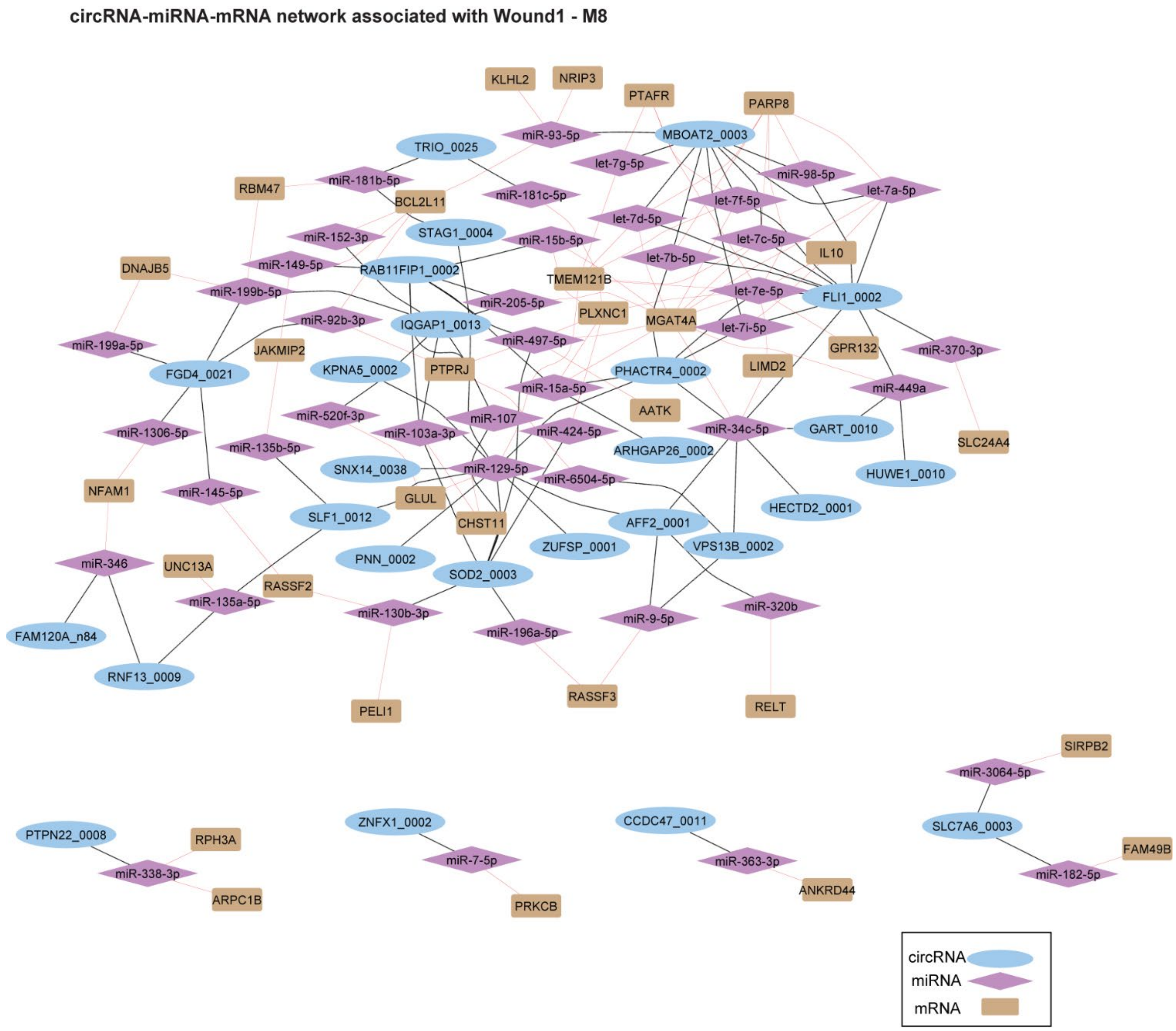


**Figure S6. CircRNA-miRNA-mRNA interactions in Wound1-associated modules.** Interaction network of the circRNAs and mRNAs that are co-expressed in the Wound1-associated module M8 and linked via miRNAs.

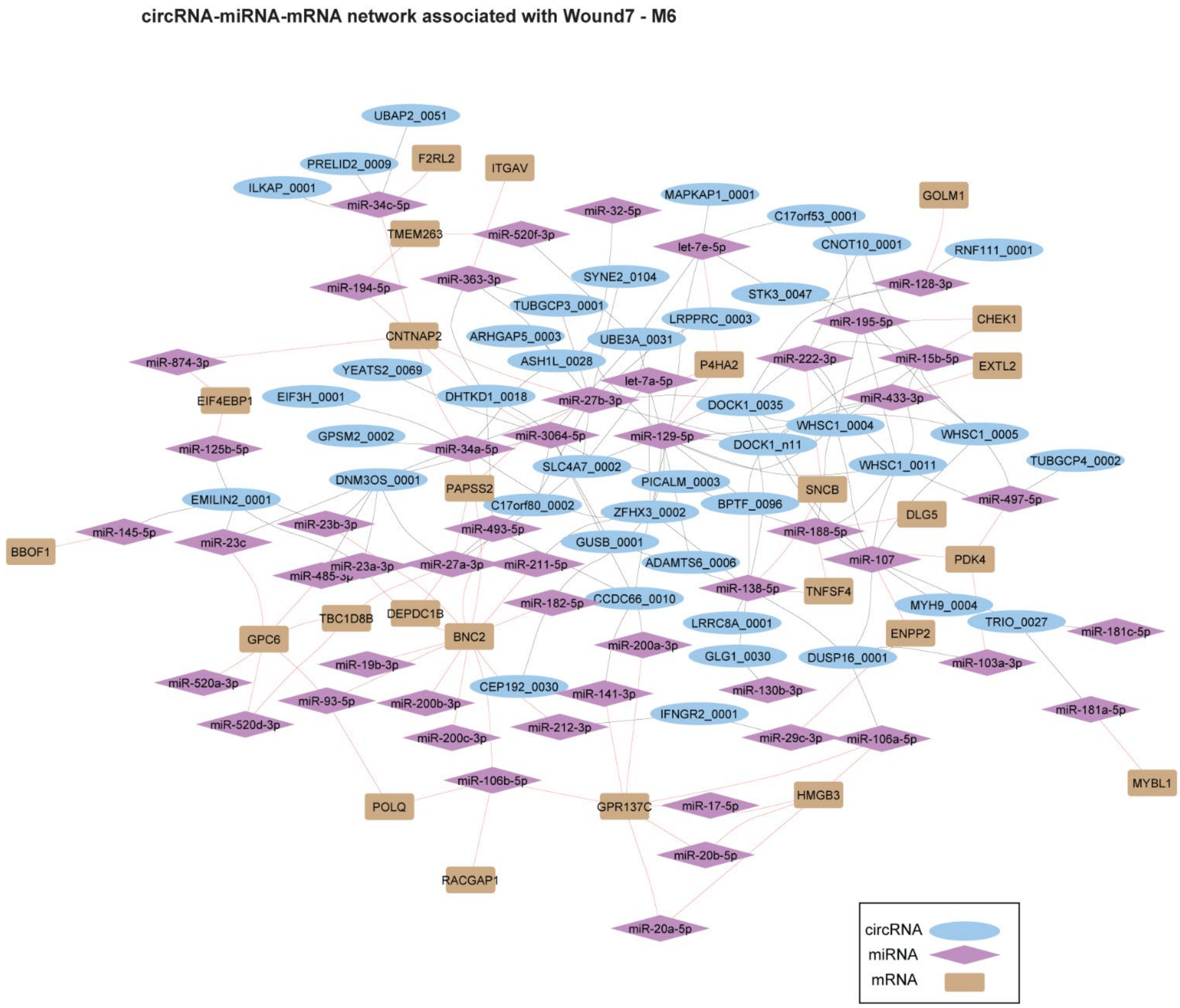


**Figure S7. CircRNA-miRNA-mRNA interactions in Wound7-associated modules.** Interaction network of the circRNAs and mRNAs that are co-expressed in the Wound7-associated module M6 and linked via miRNAs.

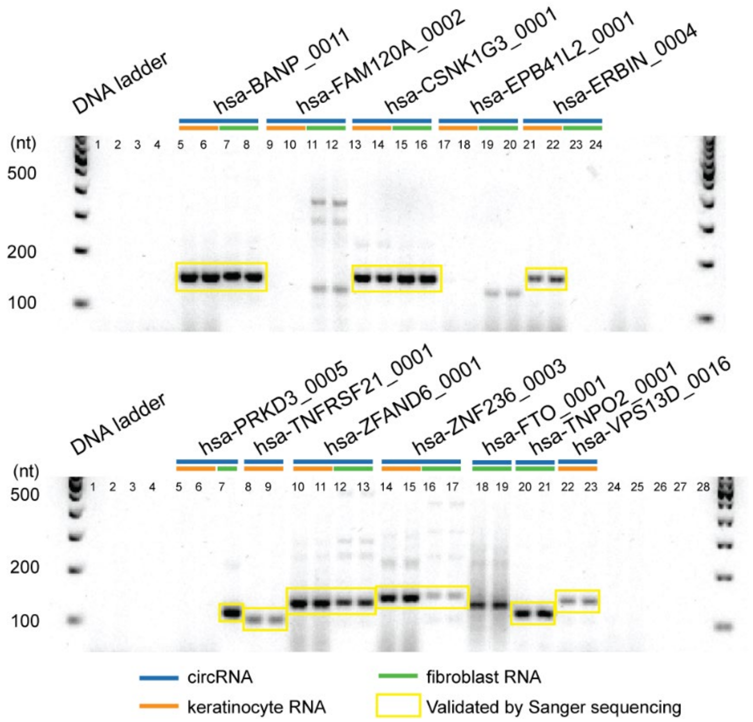


**Figure S8. Validation of circRNAs candidates.** Agarose gel electrophoresis of the PCR products on cDNA amplified with specific outward-facing primers for candidate circRNAs

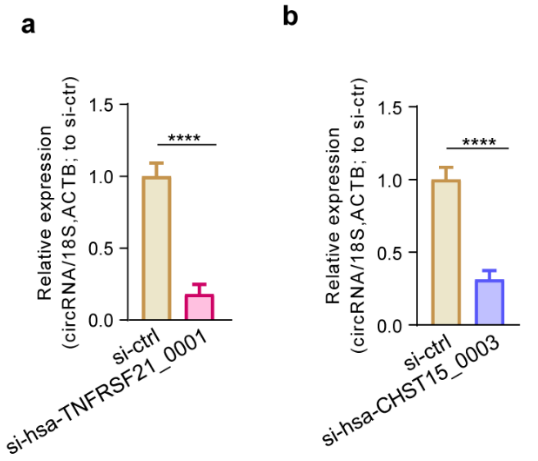


**Figure S9. siRNA knockdown efficiency.** qRT-PCR showing the expression levels of hsa-TNFRSF21_0001 (**a**) and hsa-CHST15_0003 (**b**) after 24h transfection of siRNA against hsa-TNFRSF21_0001 (**a**) and hsa-CHST15_0003 (**b**) in keratinocytes.

**Table S1**. Characteristics of Healthy Donors and Patients with VU

| Characteristics | Healthy Donors | Patients with VU |
|---|---|---|
| Study population (n) | 10 | 12 |
| Age, years (mean ± s.d.) | 65.3 ± 3.2 | 75.8 ± 12.0 |
| Ethnicity | White | White |
| Gender (male: female) | 2:8 | 5:7 |
| Biopsy location | lower leg | lower leg |
| Wound duration | Acute (1 or 7 days after injury) | 3.7 ± 5.3 years |

Abbreviations: s.d., standard deviation; VU, venous ulcer

**Table S2**. Characteristics of healthy donors

| Donor | Sex | Age | Ethnicity | Body Location | Experiment |
|---|---|---|---|---|---|
| 1 | F | 66 | caucasian | Lower leg | Long RNA sequencing |
| 2 | M | 69 | caucasian | Lower leg | Long RNA sequencing |
| 3 | F | 67 | caucasian | Lower leg | Long RNA sequencing |
| 4 | M | 69 | caucasian | Lower leg | Long RNA sequencing |
| 5 | F | 64 | caucasian | Lower leg | Long RNA sequencing |
| 6 | F | 60 | caucasian | Lower leg | qRT-PCR |
| 7 | F | 66 | caucasian | Lower leg | qRT-PCR |
| 8 | F | 60 | caucasian | Lower leg | qRT-PCR |
| 9 | F | 67 | caucasian | Lower leg | qRT-PCR |
| 10 | F | 65 | caucasian | Lower leg | qRT-PCR |

**Table S3**. Characteristics of patients with venous ulcer

| Patient | Sex | Age | Ethnicity | Wound size (cm) | Duration (years) | Body Location | Experiment |
|---|---|---|---|---|---|---|---|
| 1 | M | 86 | caucasian | 6x5 | 4 | Lower leg | Long RNA sequencing, qRT-PCR |
| 2 | F | 68 | caucasian | 15x15 | 2 | Lower leg | Long RNA sequencing, qRT-PCR |
| 3 | M | 70 | caucasian | 3x0.5 | 0.3 | Lower leg | Long RNA sequencing, qRT-PCR |
| 4 | F | 78 | caucasian | 15x12 | 1.5 | Lower leg | Long RNA sequencing, qRT-PCR |
| 5 | F | 87 | caucasian | 3x2+8x4 | 2.5 | Lower leg | Long RNA sequencing, qRT-PCR |
| 6 | F | 73 | caucasian | 3x4 | 3.5 | Lower leg | qRT-PCR |
| 7 | M | 99 | caucasian | 20x10 | 0.5 | Lower leg | qRT-PCR |
| 8 | F | 71 | caucasian | 20x20 | 20 | Lower leg | qRT-PCR |
| 9 | F | 77 | caucasian | 2.5x3+15x15 | 4.5 | Lower leg | qRT-PCR |
| 10 | M | 51 | caucasian | 2x1.5 | 3 | Lower leg | qRT-PCR |
| 11 | M | 69 | caucasian | 12x15 | 1 | Lower leg | qRT-PCR |
| 12 | F | 81 | caucasian | 7x2.5 | 1 | Lower leg | qRT-PCR |

**Table S4**. Quality control of rRNA-depleted total RNA sequencing data.
[1,2] Input reads and mapped reads are both paired fragments.

| Groups | Sample Names | Total Raw Reads | Total Clean Reads | Clean Data Rate (%) | Input reads[1] | Uniquely mapped reads[2] | Uniquely mapped rate(%) |
|---|---|---|---|---|---|---|---|
| VU | VU1 | 101622474 | 95087574 | 0.94 | 47543787 | 41697735 | 0.877 |
| VU | VU2 | 101685312 | 92173926 | 0.91 | 46086963 | 35158771 | 0.7629 |
| VU | VU3 | 132592006 | 126190658 | 0.95 | 63095329 | 55771525 | 0.8839 |
| VU | VU4 | 108786868 | 103438880 | 0.95 | 51719440 | 43012289 | 0.8316 |
| VU | VU5 | 108416676 | 101355626 | 0.93 | 50677813 | 41194500 | 0.8129 |
| Skin | Skin1 | 138065658 | 129299094 | 0.94 | 64649547 | 55300913 | 0.8554 |
| Skin | Skin2 | 102996410 | 97056760 | 0.94 | 48528380 | 38950588 | 0.8026 |
| Skin | Skin3 | 123478422 | 116018720 | 0.94 | 58009360 | 47207545 | 0.8138 |
| Skin | Skin4 | 129515874 | 115240764 | 0.89 | 57620382 | 48834112 | 0.8475 |
| Skin | Skin5 | 109239608 | 103341784 | 0.95 | 51670892 | 44434490 | 0.86 |
| Wound1 | Wound1_1 | 105127142 | 100922792 | 0.96 | 50461396 | 44012941 | 0.8722 |
| Wound1 | Wound1_2 | 104542368 | 100678896 | 0.96 | 50339448 | 44040712 | 0.8749 |
| Wound1 | Wound1_3 | 126002796 | 119469842 | 0.95 | 59734921 | 51566913 | 0.8633 |
| Wound1 | Wound1_4 | 103667154 | 98633466 | 0.95 | 49316733 | 41945529 | 0.8505 |
| Wound1 | Wound1_5 | 111446060 | 106610264 | 0.96 | 53305132 | 45076334 | 0.8456 |
| Wound7 | Wound7_1 | 115447748 | 109997276 | 0.95 | 54998638 | 45453970 | 0.8265 |
| Wound7 | Wound7_2 | 104662588 | 100033286 | 0.96 | 50016643 | 42515374 | 0.85 |
| Wound7 | Wound7_3 | 102504766 | 97652674 | 0.95 | 48826337 | 42134005 | 0.8629 |
| Wound7 | Wound7_4 | 108449990 | 103504388 | 0.95 | 51752194 | 43379024 | 0.8382 |
| Wound7 | Wound7_5 | 129466364 | 123563104 | 0.95 | 61781552 | 51511479 | 0.8338 |

**Table S5.** Differentially expressed circRNAs in human healing and nonhealing wounds

| Comparison | ID | CustomID | log2FC (circRNA) | padj (circRNA) | log2FC (mRNA) | padj (mRNA) | Type |
|---|---|---|---|---|---|---|---|
| VUvsSkin | chr1:18918362\|18919773 | hsa-IFFO2_0002 | -3.509 | 0.018 | -1.623 | 0.023 | Same_down |
| VUvsSkin | chr1:113829592\|113834439 | hsa-PTPN22_0006 | 2.519 | 0.031 | 2.370 | 0.000 | Same_up |
| VUvsSkin | chr10:7276892\|7285954 | hsa-SFMBT2_0003 | 2.304 | 0.047 | 1.532 | 0.000 | Same_up |
| VUvsSkin | chr10:95381685\|95410777 | hsa-SORBS1_0010 | -3.772 | 0.047 | -2.207 | 0.000 | Same_down |
| VUvsSkin | chr10:122082647\|122088591 | hsa-TACC2_0009 | -5.865 | 0.006 | -3.565 | 0.000 | Same_down |
| **VUvsSkin** | **chr10:124038515\|124046724** | **hsa-CHST15_0003** | **4.446** | **0.008** | **1.113** | **0.000** | **Same_up** |
| VUvsSkin | chr11:73707420\|73718890 | hsa-RAB6A_0008 | 3.738 | 0.021 | 0.440 | 0.265 | CircUp_linNot |
| VUvsSkin | chr12:55700899\|55701154 | hsa-ITGA7_0005 | -2.509 | 0.034 | -1.423 | 0.047 | Same_down |
| VUvsSkin | chr12:94169153\|94186473 | hsa-PLXNC1_0001 | 2.613 | 0.023 | 1.783 | 0.000 | Same_up |
| VUvsSkin | chr14:65561337\|65561766 | hsa-FUT8_0001 | 2.674 | 0.034 | 1.664 | 0.000 | Same_up |
| VUvsSkin | chr15:33153048\|33155045 | hsa-FMN1_0003 | -4.194 | 0.023 | -1.096 | 0.095 | CircDown_linNot |
| VUvsSkin | chr15:54012648\|54015886 | hsa-UNC13C_0040 | -4.792 | 0.002 | -4.814 | 0.000 | Same_down |
| **VUvsSkin** | **chr16:87975048\|87984259** | **hsa-BANP_0011** | **3.020** | **0.035** | **-0.227** | **0.724** | **CircUp_linNot** |
| VUvsSkin | chr17:19309885\|19329647 | hsa-EPN2_0010 | -2.637 | 0.021 | -2.220 | 0.000 | Same_down |
| VUvsSkin | chr17:39216810\|39218173 | hsa-STAC2_0002 | -5.115 | 0.018 | -9.545 | 0.000 | Same_down |
| VUvsSkin | chr19:13025021\|13025552 | hsa-NFIX_0004 | -2.922 | 0.001 | -2.142 | 0.000 | Same_down |
| VUvsSkin | chr2:131038853\|131043583 | hsa-ARHGEF4_0002 | -4.600 | 0.034 | -2.042 | 0.002 | Same_down |
| VUvsSkin | chr2:164692221\|164705105 | hsa-COBLL1_0026 | -2.479 | 0.034 | -0.204 | 0.669 | CircDown_linNot |
| VUvsSkin | chr2:205158722\|205193320 | hsa-PARD3B_0001 | -3.261 | 0.023 | -2.120 | 0.000 | Same_down |
| VUvsSkin | chr4:25847294\|25847864 | hsa-SEL1L3_0003 | 4.201 | 0.047 | 2.381 | 0.000 | Same_up |
| VUvsSkin | chr4:61934840\|62044549 | hsa-ADGRL3_0004 | -4.281 | 0.021 | -2.349 | 0.000 | Same_down |
| VUvsSkin | chr4:76738761\|76741926 | hsa-SHROOM3_0004 | -3.860 | 0.039 | -2.357 | 0.000 | Same_down |
| VUvsSkin | chr4:102304317\|102315830 | hsa-SLC39A8_0001 | 1.808 | 0.039 | 0.951 | 0.107 | CircUp_linNot |
| VUvsSkin | chr5:168488602\|168494650 | hsa-RARS_0012 | 1.447 | 0.035 | 0.936 | 0.043 | CircUp_linNot |
| **VUvsSkin** | **chr6:47283938\|47286595** | **hsa-TNFRSF21_0001** | **2.728** | **0.000** | **1.846** | **0.000** | **Same_up** |
| VUvsSkin | chr7:17868407\|17875790 | hsa-SNX13_0006 | 4.347 | 0.039 | 0.690 | 0.307 | CircUp_linNot |
| VUvsSkin | chr7:66240325\|66286709 | hsa-TPST1_0002 | 4.016 | 0.023 | 0.768 | 0.008 | CircUp_linNot |
| VUvsSkin | chr8:37870420\|37877551 | hsa-RAB11FIP1_0001 | -2.242 | 0.023 | -1.310 | 0.002 | Same_down |
| VUvsSkin | chr9:4286038\|4286523 | hsa-GLIS3_0003 | 4.025 | 0.000 | 2.375 | 0.000 | Same_up |
| **VUvsSkin** | **chr9:110972073\|110973558** | **hsa-LPAR1_0003** | **1.887** | **0.018** | **0.520** | **0.184** | **CircUp_linNot** |
| VUvsSkin | chr9:124301936\|124327445 | hsa-NEK6_0001 | 4.809 | 0.008 | 0.395 | 0.509 | CircUp_linNot |
| VUvsSkin | chr9:134690912\|134701333 | hsa-COL5A1_0001 | 2.052 | 0.034 | 1.743 | 0.000 | Same_up |
| VUvsWound1 | chr1:18918362\|18919773 | hsa-IFFO2_0002 | -2.141 | 0.007 | -0.472 | 0.379 | CircDown_linNot |
| VUvsWound1 | chr10:95381685\|95410777 | hsa-SORBS1_0010 | -4.175 | 0.018 | -1.953 | 0.000 | Same_down |
| VUvsWound1 | chr12:29293300\|29297200 | hsa-FAR2_0003 | -4.264 | 0.033 | -0.368 | 0.533 | CircDown_linNot |
| VUvsWound1 | chr2:72718103\|72733118 | hsa-EXOC6B_0002 | -1.849 | 0.012 | -0.186 | 0.626 | CircDown_linNot |
| VUvsWound1 | chr4:61934840\|62044549 | hsa-ADGRL3_0004 | -4.336 | 0.011 | -0.664 | 0.211 | CircDown_linNot |
| VUvsWound1 | chr4:102304317\|102315830 | hsa-SLC39A8_0001 | 1.995 | 0.021 | -0.490 | 0.257 | CircUp_linNot |
| VUvsWound1 | chr4:177353308\|177360677 | hsa-NEIL3_0001 | -1.874 | 0.016 | 0.109 | 0.895 | CircDown_linNot |
| VUvsWound1 | chr5:38523419\|38530666 | hsa-LIFR_0005 | -3.450 | 0.003 | -2.135 | 0.000 | Same_down |
| VUvsWound1 | chr5:83537007\|83542268 | hsa-VCAN_0003 | -1.914 | 0.000 | -0.871 | 0.004 | CircDown_linNot |
| **VUvsWound1** | **chr6:47283938\|47286595** | **hsa-TNFRSF21_0001** | **2.101** | **0.000** | **1.160** | **0.000** | **Same_up** |
| VUvsWound1 | chr9:134690912\|134701333 | hsa-COL5A1_0001 | 2.296 | 0.016 | 3.657 | 0.000 | Same_up |

| Comparison | ID | CustomID | log2FC (circRNA) | padj (circRNA) | log2FC (mRNA) | padj (mRNA) | Type |
|---|---|---|---|---|---|---|---|
| VUvsWound1 | chrX:148661908\|148662768 | hsa-AFF2_0002 | -2.695 | 0.000 | -0.095 | 0.861 | CircDown_linNot |
| VUvsWound7 | chr1:18918362\|18919773 | hsa-IFFO2_0002 | -2.123 | 0.002 | -0.371 | 0.569 | CircDown_linNot |
| VUvsWound7 | chr10:126970702\|127127764 | hsa-DOCK1_0035 | -1.714 | 0.000 | -0.050 | 0.898 | CircDown_linNot |
| VUvsWound7 | chr4:105233897\|105237351 | hsa-TET2_0005 | -1.162 | 0.048 | -0.001 | 0.998 | CircDown_linNot |
| VUvsWound7 | chr5:83537007\|83542268 | hsa-VCAN_0003 | -1.208 | 0.016 | -0.612 | 0.045 | CircDown_linNot |
| VUvsWound7 | chr5:95755396\|95763620 | hsa-RHOBTB3_0011 | 1.934 | 0.002 | 0.447 | 0.128 | CircUp_linNot |
| **VUvsWound7** | **chr9:110972073\|110973558** | **hsa-LPAR1_0003** | **1.261** | **0.011** | **0.246** | **0.588** | **CircUp_linNot** |
| Wound7vsSkin | chr1:22078429\|22086866 | hsa-CDC42_0001 | 4.313 | 0.024 | 0.966 | 0.000 | CircUp_linNot |
| Wound7vsSkin | chr1:44411981\|44412722 | hsa-RNF220_0002 | -2.474 | 0.021 | -3.601 | 0.000 | Same_down |
| **Wound7vsSkin** | **chr1:155438327\|155459898** | **hsa-ASH1L_0028** | **2.434** | **0.026** | **-0.201** | **0.354** | **CircUp_linNot** |
| Wound7vsSkin | chr10:126970702\|127127764 | hsa-DOCK1_0035 | 1.274 | 0.042 | 0.115 | 0.540 | CircUp_linNot |
| Wound7vsSkin | chr12:55700899\|55701154 | hsa-ITGA7_0005 | -3.279 | 0.007 | -2.402 | 0.000 | Same_down |
| Wound7vsSkin | chr12:121416159\|121417911 | hsa-RNF34_0005 | -3.981 | 0.046 | -0.446 | 0.131 | CircDown_linNot |
| Wound7vsSkin | chr12:123498544\|123499536 | hsa-RILPL1_0001 | -2.574 | 0.026 | -1.461 | 0.001 | Same_down |
| Wound7vsSkin | chr15:42665253\|42669337 | hsa-STARD9_0011 | -3.653 | 0.037 | -2.369 | 0.000 | Same_down |
| Wound7vsSkin | chr15:54012648\|54015886 | hsa-UNC13C_0040 | -2.753 | 0.015 | -3.033 | 0.000 | Same_down |
| Wound7vsSkin | chr16:53873786\|53934109 | hsa-FTO_0003 | -3.599 | 0.037 | -0.918 | 0.000 | CircDown_linNot |
| Wound7vsSkin | chr16:70463169\|70463774 | hsa-FUK_0007 | -3.557 | 0.026 | -2.091 | 0.000 | Same_down |
| Wound7vsSkin | chr16:84740309\|84745673 | hsa-USP10_0002 | 2.391 | 0.041 | 0.450 | 0.055 | CircUp_linNot |
| Wound7vsSkin | chr17:39216810\|39218173 | hsa-STAC2_0002 | -4.194 | 0.015 | -3.551 | 0.000 | Same_down |
| Wound7vsSkin | chr17:40495987\|40496520 | hsa-TNS4_0001 | -2.686 | 0.026 | -0.623 | 0.159 | CircDown_linNot |
| Wound7vsSkin | chr18:12999421\|13042334 | hsa-CEP192_0030 | 3.661 | 0.042 | 0.320 | 0.283 | CircUp_linNot |
| Wound7vsSkin | chr18:44949881\|44953340 | hsa-SETBP1_0001 | -2.340 | 0.015 | -1.012 | 0.000 | Same_down |
| **Wound7vsSkin** | **chr18:76849526\|76851939** | **hsa-ZNF236_0003** | **2.070** | **0.027** | **-0.242** | **0.346** | **CircUp_linNot** |
| Wound7vsSkin | chr19:3623688\|3624162 | hsa-CACTIN_0003 | -2.505 | 0.026 | -1.438 | 0.009 | Same_down |
| Wound7vsSkin | chr19:47264603\|47264946 | hsa-CCDC9_0002 | -3.071 | 0.020 | -2.449 | 0.002 | Same_down |
| Wound7vsSkin | chr2:61522611\|61533903 | hsa-XPO1_0001 | 1.666 | 0.026 | 1.027 | 0.007 | Same_up |
| Wound7vsSkin | chr2:205158722\|205193320 | hsa-PARD3B_0001 | -2.143 | 0.041 | -2.440 | 0.000 | Same_down |
| Wound7vsSkin | chr21:15762891\|15766141 | hsa-USP25_0001 | 4.745 | 0.007 | -0.083 | 0.658 | CircUp_linNot |
| **Wound7vsSkin** | **chr22:36341370\|36349255** | **hsa-MYH9_0004** | **4.085** | **0.026** | **1.183** | **0.000** | **Same_up** |
| Wound7vsSkin | chr3:27411642\|27452498 | hsa-SLC4A7_0002 | 3.730 | 0.034 | 0.740 | 0.000 | CircUp_linNot |
| Wound7vsSkin | chr3:136980466\|136995556 | hsa-IL20RB_0001 | -1.732 | 0.041 | -1.944 | 0.000 | Same_down |
| Wound7vsSkin | chr3:149846011\|149921227 | hsa-RNF13_0008 | 1.479 | 0.037 | 0.577 | 0.099 | CircUp_linNot |
| Wound7vsSkin | chr4:76134175\|76144473 | hsa-NUP54_0020 | 2.586 | 0.020 | 0.732 | 0.010 | CircUp_linNot |
| Wound7vsSkin | chr4:76738761\|76741926 | hsa-SHROOM3_0004 | -3.561 | 0.041 | -2.518 | 0.000 | Same_down |
| Wound7vsSkin | chr4:105233897\|105237351 | hsa-TET2_0005 | 1.544 | 0.026 | -0.128 | 0.537 | CircUp_linNot |
| Wound7vsSkin | chr4:177353308\|177360677 | hsa-NEIL3_0001 | 2.466 | 0.007 | 1.273 | 0.009 | Same_up |
| Wound7vsSkin | chr5:65451475\|65473952 | hsa-ADAMTS6_0006 | 4.624 | 0.006 | 3.088 | 0.000 | Same_up |
| **Wound7vsSkin** | **chr5:66053406\|66054951** | **hsa-ERBIN_0004** | **1.728** | **0.037** | **0.844** | **0.098** | **CircUp_linNot** |
| Wound7vsSkin | chr5:69174877\|69175537 | hsa-CCNB1_0001 | 4.126 | 0.020 | 2.857 | 0.000 | Same_up |
| **Wound7vsSkin** | **chr5:123545417\|123557564** | **hsa-CSNK1G3_0001** | **1.365** | **0.026** | **0.625** | **0.022** | **CircUp_linNot** |
| Wound7vsSkin | chr5:168488602\|168494650 | hsa-RARS_0012 | 1.313 | 0.036 | 1.421 | 0.000 | Same_up |
| Wound7vsSkin | chr6:13579451\|13584225 | hsa-SIRT5_0007 | 3.705 | 0.033 | -0.895 | 0.002 | CircUp_linNot |
| **Wound7vsSkin** | **chr6:47283938\|47286595** | **hsa-TNFRSF21_0001** | **1.887** | **0.002** | **1.076** | **0.005** | **Same_up** |
| **Wound7vsSkin** | **chr6:130926605\|130956499** | **hsa-EPB41L2_0001** | **-2.665** | **0.008** | **-0.188** | **0.458** | **CircDown_linNot** |

| Comparison | ID | CustomID | log2FC (circRNA) | padj (circRNA) | log2FC (linRNA) | padj (linRNA) | Type |
|---|---|---|---|---|---|---|---|
| Wound7vsSkin | chr7:17868407\|17875790 | hsa-SNX13_0006 | 4.342 | 0.024 | 0.550 | 0.227 | CircUp_linNot |
| Wound7vsSkin | chr7:98191614\|98194572 | hsa-LMTK2_n76 | -1.888 | 0.037 | -1.445 | 0.002 | Same_down |
| Wound7vsSkin | chr8:37870420\|37877551 | hsa-RAB11FIP1_0001 | -2.641 | 0.007 | -1.098 | 0.003 | Same_down |
| Wound7vsSkin | chr8:38429682\|38429948 | hsa-FGFR1_0001 | -2.455 | 0.002 | -1.987 | 0.000 | Same_down |
| Wound7vsSkin | chr8:121628714\|121629340 | hsa-HAS2_0001 | 2.495 | 0.041 | 2.706 | 0.000 | Same_up |
| Wound7vsSkin | chr9:4286038\|4286523 | hsa-GLIS3_0003 | 4.048 | 0.000 | 2.122 | 0.000 | Same_up |
| Wound7vsSkin | chr9:37424845\|37426654 | hsa-GRHPR_0002 | -1.450 | 0.024 | -1.728 | 0.002 | Same_down |
| Wound7vsSkin | chr9:111386377\|111391824 | hsa-KIAA0368_0046 | 2.208 | 0.005 | 0.343 | 0.042 | CircUp_linNot |
| Wound7vsSkin | chr9:134690912\|134701333 | hsa-COL5A1_0001 | 1.923 | 0.037 | 1.349 | 0.003 | Same_up |
| Wound7vsSkin | chr9:134824600\|134825904 | hsa-COL5A1_0005 | 4.022 | 0.026 | 0.764 | 0.190 | CircUp_linNot |
| Wound1vsSkin | chr1:16201782\|16202574 | hsa-ARHGEF19_0001 | -1.964 | 0.048 | -2.082 | 0.001 | Same_down |
| Wound1vsSkin | chr1:20770930\|20773610 | hsa-HP1BP3_0016 | 1.773 | 0.049 | -0.082 | 0.601 | CircUp_linNot |
| Wound1vsSkin | chr1:113829592\|113834439 | hsa-PTPN22_0006 | 3.301 | 0.000 | 1.585 | 0.004 | Same_up |
| Wound1vsSkin | chr10:101667886\|101676436 | hsa-FBXW4_0001 | -1.728 | 0.019 | -2.216 | 0.000 | Same_down |
| Wound1vsSkin | chr10:122082647\|122088591 | hsa-TACC2_0009 | -3.755 | 0.029 | -3.439 | 0.000 | Same_down |
| Wound1vsSkin | chr11:86007542\|86031611 | hsa-PICALM_0007 | 2.403 | 0.049 | 1.394 | 0.000 | Same_up |
| Wound1vsSkin | chr12:29293300\|29297200 | hsa-FAR2_0003 | 4.234 | 0.013 | -0.211 | 0.816 | CircUp_linNot |
| Wound1vsSkin | chr12:94169153\|94186473 | hsa-PLXNC1_0001 | 2.440 | 0.027 | 1.254 | 0.000 | Same_up |
| Wound1vsSkin | chr12:123498544\|123499536 | hsa-RILPL1_0001 | -2.008 | 0.049 | -1.347 | 0.004 | Same_down |
| Wound1vsSkin | chr15:33153048\|33155045 | hsa-FMN1_0003 | -3.996 | 0.019 | -2.096 | 0.000 | Same_down |
| **Wound1vsSkin** | **chr15:80120328\|80122800** | **hsa-ZFAND6_0001** | **2.253** | **0.003** | **0.629** | **0.001** | **CircUp_linNot** |
| Wound1vsSkin | chr16:4466153\|4469465 | hsa-NMRAL1_0001 | -2.274 | 0.023 | -2.030 | 0.002 | Same_down |
| Wound1vsSkin | chr16:31361836\|31362753 | hsa-ITGAX_0001 | 4.828 | 0.003 | 4.105 | 0.000 | Same_up |
| Wound1vsSkin | chr16:84740309\|84745673 | hsa-USP10_0002 | 2.497 | 0.029 | 0.841 | 0.000 | CircUp_linNot |
| Wound1vsSkin | chr17:19309885\|19329647 | hsa-EPN2_0010 | -1.818 | 0.029 | -1.954 | 0.000 | Same_down |
| Wound1vsSkin | chr18:44701175\|44701832 | hsa-SETBP1_0004 | -1.417 | 0.046 | -1.618 | 0.000 | Same_down |
| Wound1vsSkin | chr19:45398005\|45398339 | hsa-PPP1R13L_0002 | -4.372 | 0.005 | -2.860 | 0.000 | Same_down |
| Wound1vsSkin | chr2:8908621\|8958642 | hsa-MBOAT2_0003 | 3.096 | 0.000 | -0.569 | 0.120 | CircUp_linNot |
| **Wound1vsSkin** | **chr2:37316237\|37317179** | **hsa-PRKD3_0005** | **1.453** | **0.029** | **0.096** | **0.827** | **CircUp_linNot** |
| Wound1vsSkin | chr2:61522611\|61533903 | hsa-XPO1_0001 | 2.103 | 0.001 | 1.066 | 0.006 | Same_up |
| Wound1vsSkin | chr21:15762891\|15766141 | hsa-USP25_0001 | 4.186 | 0.029 | 0.131 | 0.487 | CircUp_linNot |
| Wound1vsSkin | chr21:29326059\|29329693 | hsa-BACH1_0002 | 2.026 | 0.022 | 0.463 | 0.025 | CircUp_linNot |
| Wound1vsSkin | chr4:87195324\|87195690 | hsa-KLHL8_0015 | 1.873 | 0.002 | 0.953 | 0.008 | CircUp_linNot |
| Wound1vsSkin | chr4:139137630\|139139497 | hsa-ELF2_0004 | 3.086 | 0.011 | 0.250 | 0.213 | CircUp_linNot |
| Wound1vsSkin | chr4:177353308\|177360677 | hsa-NEIL3_0001 | 3.216 | 0.000 | 0.681 | 0.251 | CircUp_linNot |
| **Wound1vsSkin** | **chr5:66053406\|66054951** | **hsa-ERBIN_0004** | **1.720** | **0.019** | **0.496** | **0.387** | **CircUp_linNot** |
| Wound1vsSkin | chr5:83537007\|83542268 | hsa-VCAN_0003 | 1.517 | 0.012 | 1.834 | 0.000 | Same_up |
| Wound1vsSkin | chr5:83537007\|83555038 | hsa-VCAN_0001 | 2.537 | 0.044 | 1.513 | 0.002 | Same_up |
| **Wound1vsSkin** | **chr5:123545417\|123557564** | **hsa-CSNK1G3_0001** | **1.380** | **0.024** | **0.399** | **0.210** | **CircUp_linNot** |
| Wound1vsSkin | chr5:137985257\|137988315 | hsa-FAM13B_0019 | 1.791 | 0.001 | 1.041 | 0.000 | Same_up |
| Wound1vsSkin | chr5:143054439\|143057747 | hsa-ARHGAP26_0002 | 2.208 | 0.004 | 1.201 | 0.000 | Same_up |
| Wound1vsSkin | chr5:168488602\|168494650 | hsa-RARS_0012 | 1.490 | 0.011 | 1.421 | 0.000 | Same_up |
| Wound1vsSkin | chr5:177536358\|177539147 | hsa-FAM193B_0003 | -1.819 | 0.019 | -1.948 | 0.001 | Same_down |
| Wound1vsSkin | chr7:17868407\|17875790 | hsa-SNX13_0006 | 4.588 | 0.008 | 0.856 | 0.069 | CircUp_linNot |
| Wound1vsSkin | chr7:32632543\|32639365 | hsa-DPY19L1P1_0004 | 4.237 | 0.048 | 1.177 | 0.036 | Same_up |

| Comparison | ID | CustomID | log2FC (circRNA) | padj (circRNA) | log2FC (mRNA) | padj (mRNA) | Type |
|---|---|---|---|---|---|---|---|
| Wound1vsSkin | chr7:98191614\|98194572 | hsa-LMTK2_n76 | -2.002 | 0.020 | -1.145 | 0.020 | Same_down |
| Wound1vsSkin | chr7:134947508\|134950514 | hsa-CALD1_0004 | -3.862 | 0.037 | -0.818 | 0.031 | CircDown_linNot |
| Wound1vsSkin | chr8:37870420\|37877551 | hsa-RAB11FIP1_0001 | -2.148 | 0.011 | -0.716 | 0.079 | CircDown_linNot |
| Wound1vsSkin | chr8:38819522\|38820635 | hsa-TACC1_0004 | 4.153 | 0.048 | -0.216 | 0.476 | CircUp_linNot |
| Wound1vsSkin | chr9:4286038\|4286523 | hsa-GLIS3_0003 | 3.337 | 0.005 | 2.638 | 0.000 | Same_up |
| Wound1vsSkin | chr9:93471141\|93476338 | hsa-FAM120A_0002 | 2.268 | 0.026 | 0.417 | 0.009 | CircUp_linNot |
| Wound1vsSkin | chrX:148661908\|148662768 | hsa-AFF2_0002 | 1.402 | 0.013 | -0.970 | 0.006 | CircUp_linNot |
| Wound7vsWound1 | chr1:113829592\|113834439 | hsa-PTPN22_0006 | -2.039 | 0.010 | -0.064 | 0.928 | CircDown_linNot |
| Wound7vsWound1 | chr10:126970702\|127127764 | hsa-DOCK1_0035 | 1.430 | 0.017 | 0.301 | 0.100 | CircUp_linNot |
| Wound7vsWound1 | chr2:8908621\|8958642 | hsa-MBOAT2_0003 | -1.306 | 0.017 | 0.854 | 0.000 | CircDown_linNot |
| Wound7vsWound1 | chr5:38523419\|38530666 | hsa-LIFR_0005 | -2.312 | 0.009 | -0.862 | 0.000 | CircDown_linNot |
| **Wound7vsWound1** | **chr6:47283938\|47286595** | **hsa-TNFRSF21_0001** | **1.336** | **0.011** | **0.379** | **0.204** | **CircUp_linNot** |
| Wound7vsWound1 | chr9:134690912\|134701333 | hsa-COL5A1_0001 | 2.237 | 0.012 | 3.252 | 0.000 | Same_up |
| Wound7vsWound1 | chrX:148661908\|148662768 | hsa-AFF2_0002 | -1.931 | 0.000 | -0.927 | 0.028 | CircDown_linNot |

**Table S6.** Primers for circRNA detection

| ID | CustomID | Forward primer | Reverse primer | PCR product size (bp) |
|---|---|---|---|---|
| chr16:87975048\|87984259 | hsa-BANP_0011 | GATCTGGTCACGAACAAGCA | CCGGTCAACACGAAAGAGTT | 145 |
| chr6:47283938\|47286595 | hsa-TNFRSF21_0001 | TGAGCTGTGGTGGTGCTAAG | AAAGGCAGGGCTGAAGAAAT | 103 |
| chr10:124038515\|124046724 | hsa-CHST15_0003 | TGTGGGGCGCAAACTTTAAA | CTACGACAACAGCACGGATG | 142 |
| chr9:110972073\|110973558 | hsa-LPAR1_0003 | AGCTGTGTACCTGATGCTGT | GGCTGCCATCTCTACTTCCA | 131 |
| chr5:123545417\|123557564 | hsa-CSNK1G3_0001 | GCACCACAGCTACATTTGGA | GGAGCATGTTCATCCCATTC | 155 |
| chr5:66053406\|66054951 | hsa-ERBIN_0004 | CAAATGCAAGGGTTGGTTCT | TCTCCATTTTGTAAAAGGCTGTT | 158 |
| chr2:37316237\|37317179 | hsa-PRKD3_0005 | AGATGAATGGGTCCATCGAG | TACCATTGAAGCCCAGGAAC | 121 |
| chr15:80120328\|80122800 | hsa-ZFAND6_0001 | TGTGGACAAAGCAGTACCTGA | CACATGCCATTTGTACGAGG | 144 |
| chr6:130926605\|130956499 | hsa-EPB41L2_0001 | TCAGTAGTCATGGCCACAGC | AAACATGCCAAGGGACAAGT | 130 |
| chr18:76849526\|76851939 | hsa-ZNF236_0003 | AGTGGCTAGTCTCAAAGCGC | GCGTTGAAACTGGGATTCTT | 159 |
| chr22:36341370\|36349255 | hsa-MYH9_0004 | GCCAGCGGATTGTTGATGAA | GCAAGAAGAGGCACGAGATG | 131 |
| chr1:155438327\|155459898 | hsa-ASH1L_0028 | GAAGACCTTTTCCGGGTAGG | GCAGAAGCCATTACCAGAGG | 124 |
| chr19:12714821\|12715739 | hsa-TNPO2_0001 | CGATGGTGGTGATGAGAATG | ACCCAGGACCATGATGAGAA | 125 |
| chr1:12275825\|12278038 | hsa-VPS13D_0016 | CTGTTGGGTCTGAAGGAAGC | CCCATCCCTACTTTTCCTCC | 149 |
